# TMEM135 deficiency remodels hepatic lipid homeostasis and protein malonylation through a DHA-sensitive lipogenic program buffered by peroxisomal metabolism

**DOI:** 10.64898/2026.07.22.740069

**Authors:** Ryo Hagimori, Michael Landowski, Siyu Song, Purnima Gogoi, Richard S. Brush, Sarah M. Bonvicino, Greg Barrett-Wilt, Sakae Ikeda, Ken-ichi Yamada, Chi-Liang Eric Yen, Tetsuya Takimoto, Martin-Paul Agbaga, Akihiro Ikeda

## Abstract

TMEM135 has been implicated in lipid metabolism, but its role in regulating hepatic lipid homeostasis remains unclear. Here, we investigated how TMEM135 affects hepatic lipid metabolism using *Tmem135* mutant mice with liver-specific *Pex5* deletion. The *Tmem135* mutation induced a lipogenic state characterized by depletion of docosahexaenoic acid (DHA), activation of SREBP-dependent pathways, and increased monounsaturated fatty acids without causing hepatic steatosis. In contrast, loss of PEX5-dependent peroxisomal function in *Tmem135* mutant mice resulted in marked hepatic lipid accumulation, indicating that peroxisomal metabolism buffers the elevated lipogenic state. Fish oil supplementation to *Tmem135* mutant mice restored DHA levels and suppressed lipogenesis. Proteomics identified distinct DHA-sensitive metabolic programs, including activation of SREBP-dependent lipogenesis. Quantitative malonyl-proteomics revealed increased malonylation of glycolytic enzymes, accompanied by altered glycolytic output. Together, these findings identify TMEM135 as a central regulator of hepatic lipid metabolism and uncover a coordinated mechanism linking lipid availability, lipogenesis, and post-translational metabolic regulation.

**Highlight:** *Tmem135* mutation induces a lipogenic state with increased lipolysis without hepatic steatosis. PEX5-dependent peroxisomal function buffers lipid accumulation in *Tmem135* mutant liver.

Fish oil supplementation restores DHA and suppresses SREBP-dependent lipogenesis in *Tmem135* mutants.

DHA-sensitive protein malonylation targets glycolytic enzymes in *Tmem135* mutant liver.

## INTRODUCTION

The liver plays a central role in maintaining systemic lipid homeostasis by coordinating lipid synthesis, oxidation, storage, and secretion^1,2^. Disruption of this balance can lead to hepatic steatosis, a pathological accumulation of lipids in hepatocytes that underlies metabolic disorders such as non-alcoholic fatty liver disease (NAFLD)^3^. Hepatic lipid levels are determined not only by lipid synthesis rates but also by the capacity of metabolic pathways that process, partition, and redistribute lipid intermediates. Understanding how lipid homeostasis is regulated and buffered under conditions of increased lipid synthesis remains an important challenge in metabolic biology.

The peroxisome-associated transmembrane protein 135 (TMEM135) is an emerging regulator of lipid metabolism, particularly docosahexaenoic acid (22:6n-3; DHA), that plays important roles in cellular function^4–8^, yet the molecular mechanisms underlying its role in hepatic lipid homeostasis remain incompletely defined. Emerging evidence suggests that TMEM135 influences lipid metabolic pathways, including processes linked to fatty acid synthesis and remodeling^7^, and may impact the balance between lipid production, utilization, and export. Notably, the *Tmem135* mutation has been reported to ameliorate hepatic steatosis in metabolic disease models^4,8^, including *ob/ob* mice, where activation of PPARα-associated fatty acid oxidation pathways is implicated in this protective effect^8^. In addition, the *Tmem135* mutation has been associated with an increase in peroxisome abundance, suggesting a potential link between TMEM135 function and peroxisomal metabolic capacity^8^. Furthermore, lipid availability—particularly polyunsaturated fatty acids (PUFA) such as DHA—is known to regulate lipogenic programs through SREBP signaling ^9–11^, suggesting that TMEM135 may influence lipid metabolism by modulating lipid composition and downstream regulatory pathways. Beyond lipogenic regulation, hepatic lipid homeostasis both reflects and interacts with transcriptional programs that maintain overall hepatocellular function^12–15^, raising the possibility that TMEM135-dependent remodeling may affect broader aspects of hepatic metabolism beyond lipid synthesis.

Peroxisomes are key organelles involved in lipid metabolism, particularly in the processing of very-long-chain fatty acids and PUFA via β-oxidation pathways^16–18^. In addition to fatty acid oxidation, peroxisomes participate in lipid remodeling and metabolic interactions with mitochondria and other cellular compartments^17,19^. TMEM135 has been shown to link peroxisomal function to mitochondrial dynamics and energy homeostasis^6^. While defects in peroxisomal function are known to disrupt lipid metabolism^16^, how peroxisomal capacity contributes to maintaining lipid homeostasis under conditions of altered lipid handling remains incompletely understood. Particularly, it is unclear whether peroxisomes play an active role in buffering lipid metabolic stress arising from dysregulated lipid metabolism, including that associated with TMEM135-dependent metabolic remodeling.

Here we investigated how the *Tmem135* mutation alters hepatic lipid homeostasis, and whether peroxisomal function contributes to this process. To address this, we combined a mouse carrying the *Tmem135* mutation with liver-specific disruption of peroxisomal protein import through deletion of *Pex5*, allowing us to directly test the contribution of peroxisomal metabolic capacity to TMEM135-dependent lipid remodeling. Our results reveal that the *Tmem135* mutation induces a lipogenic metabolic state that is normally buffered by peroxisomal function, and that loss of PEX5-dependent peroxisomal activity uncovers this latent metabolic imbalance, leading to hepatic lipid accumulation.

We further investigated the mechanisms underlying *Tmem135* mutation induced lipogenic state by examining lipid availability and SREBP-dependent regulation. Using integrated lipidomic and proteomic analyses, we demonstrate that the *Tmem135* mutation promotes increased production of monounsaturated fatty acids (MUFA), including 18:1, through upregulation of SCD1 and activation of a lipogenic program that is selectively suppressed by fish oil supplementation (DHA-rich diet). Finally, we show that this lipogenic state is associated with increased protein malonylation, revealing a post-translational layer of metabolic regulation.

Together, these findings identify TMEM135 as a central regulator of hepatic lipid metabolism and demonstrate that peroxisomal function serves as a buffering system that determines whether a lipogenic state is maintained as balanced lipid homeostasis or diverted toward lipid storage. In addition, our results reveal that lipid availability and post-translational modification cooperate to coordinate broader metabolic adaptation downstream of TMEM135-dependent remodeling.

## RESULTS

### Loss of peroxisomal function exacerbates lipid accumulation in *Tmem135* mutant liver

The *Tmem135* mutation has been shown to ameliorate hepatic steatosis in both *ob*/*ob* and high-fat diet–induced models of NAFLD^4,8^, suggesting a role in regulating hepatic lipid metabolism. Although the underlying mechanisms remain incompletely understood, alterations in lipid handling pathways have been implicated. Given emerging evidence linking TMEM135 in peroxisomal function^5,8^, and the central role of peroxisomes in fatty acid metabolism^17,20^, we hypothesized that TMEM135-dependent lipid remodeling generates a metabolic state that is buffered by peroxisomal function. To test this, we crossed *Tmem135^FUN025/FUN025^* (*Tmem135* mutant) mice with liver-specific *Pex5* knockout mice (*Alb-Cre*;*Pex5^fl/fl^*) to generate *Alb-Cre*;*Pex5^fl/fl^*;*Tmem135^FUN025/FUN025^* mice in which hepatic peroxisomal protein import and peroxisomal metabolism are disrupted. We first confirmed efficient depletion of PEX5 in the livers of *Alb-Cre*;*Pex5^fl/fl^* and *Alb-Cre*;*Pex5^fl/fl^*;*Tmem135^FUN025/FUN025^* mice by immunoblot analysis (Figure S1A). Consistent with impaired peroxisomal maintenance, both immunoblotting and immunohistochemistry for the peroxisomal membrane proteins PMP70 and PEX14 demonstrated a marked reduction in hepatic peroxisomal abundance in *Pex5*-deficient livers (Figure S1B, S1C, and S1D).

We examined hepatic lipid accumulation in control, *Tmem135* mutant, liver-specific *Pex5* knockout, and liver-specific *Pex5*-deficient *Tmem135* mutant mice. Histological analysis by H&E staining revealed that control, *Tmem135* mutant, and *Pex5*-deficient livers exhibited largely preserved hepatic architecture without overt steatosis, whereas the liver-specific *Pex5*-deficient *Tmem135* mutant livers displayed prominent accumulation of large, clear cytoplasmic vacuoles consistent with lipid droplets, indicative of marked hepatic steatosis (Figure 1A). Consistent with this, BODIPY staining showed relatively limited lipid droplet accumulation in control, *Tmem135* mutant, and *Pex5*-deficient livers, while the double mutant mice exhibited a striking increase in lipid droplet abundance, confirming that substantial neutral lipid storage emerges primarily when the *Tmem135* mutation is combined with loss of peroxisomal function (Figure 1B, C). At higher resolution, however, *Tmem135* mutant hepatocytes appeared to contain relatively larger lipid droplets compared with controls, despite the absence of overt steatosis, raising the possibility that the TMEM135 mutation alters lipid droplet morphology before causing major bulk lipid accumulation. This effect appeared more pronounced in the double mutant, which showed both increased lipid droplet abundance and enlarged droplet structures (Figure 1D).

**Figure 1.**
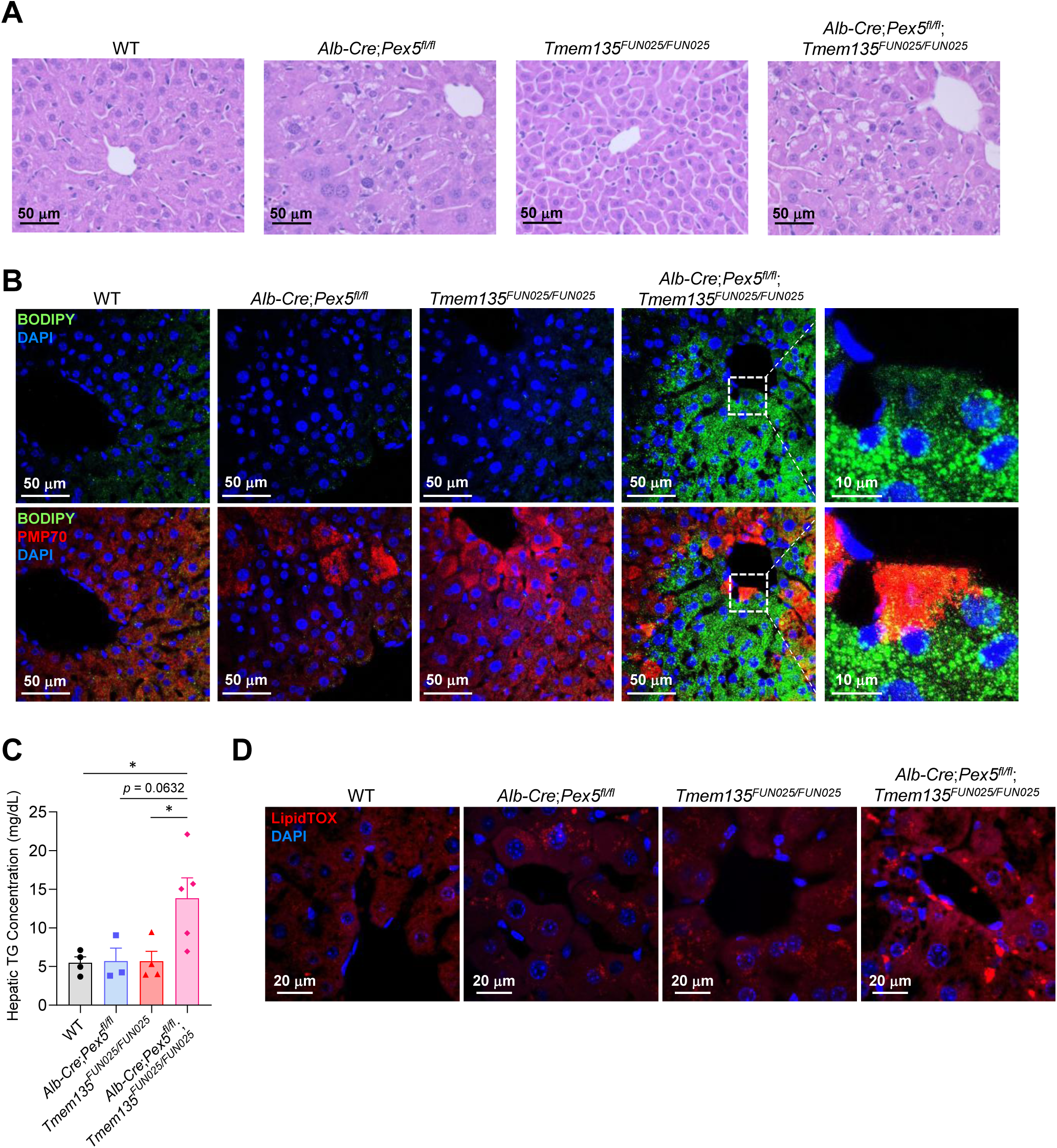
Peroxisomal dysfunction unmasks hepatic steatosis in Tmem135 mutant mice. (A) Representative images of hematoxylin and eosin (H&E) stained liver sections from 3-month WT, *Alb-Cre*;*Pex5^fl/fl^*, *Tmem135^FUN025/FUN025^*, and *Alb-Cre*;*Pex5^fl/fl^*;*Tmem135^FUN025/FUN025^* mice. Scale bar, 50 μm. (B) Representative fluorescence images showing BODIPY (green), PEX70 immunostaining (red), and DAPI (blue) in livers from 3-month-old WT, *Alb-Cre*;*Pex5^fl/fl^*, *Tmem135^FUN025/FUN025^*, and *Alb-Cre*;*Pex5^fl/fl^*;*Tmem135^FUN025/FUN025^* mice. White boxes indicate regions shown at higher magnification in the *Alb-Cre*;*Pex5^fl/fl^*;*Tmem135^FUN025/FUN025^* livers. Scale bars, 50 μm (main images) and 10 μm (enlarged images). (C) Quantification of hepatic triglyceride (TG) concentrations in livers from 3-month-old WT, *Alb-Cre*;*Pex5^fl/fl^*, *Tmem135^FUN025/FUN025^*, and *Alb-Cre*;*Pex5^fl/fl^*;*Tmem135^FUN025/FUN025^* mice. Data are presented as mean ± SEM (n = 3–5 mice per group). Statistical significance was determined by two-way ANOVA followed by Šídák’s multiple-comparisons test. \**p* < 0.05 (D) Representative high-resolution confocal micrographs of LipidTOX (red) and DAPI (blue) staining in livers from 3-month-old WT, *Alb-Cre*;*Pex5^fl/fl^*, *Tmem135^FUN025/FUN025^*, and *Alb-Cre*;*Pex5^fl/fl^*;*Tmem135^FUN025/FUN025^* mice. Scale bar, 20 μm.

These results suggest that the *Tmem135* mutation may establish a lipid-handling state associated with altered droplet morphology, while intact peroxisomal function prevents progression to overt steatosis; when peroxisomal capacity is disrupted, this buffered state collapses into marked hepatic lipid accumulation.

### *Tmem135* mutation promotes MUFA-enriched lipid remodeling without inducing steatosis

To define the lipid changes underlying these phenotypes, we performed lipidomic analysis of liver and plasma. Analysis of total fatty acid composition revealed a significant increase in overall fatty acid levels in *Pex5*-deficient and double-mutant livers, whereas levels in *Tmem135* mutant mice were comparable to wild-type (Figure 2A). In contrast, 22:6n-3 (DHA) was markedly reduced in *Tmem135* mutant and double-mutant livers, and significantly decreased, though still detectable, in *Pex5*-deficient liver. In plasma, total fatty acid levels were not significantly altered across genotypes, although 22:6 showed a similar decreasing trend (Figure 2B). Together, these data indicate that loss of peroxisomal function leads to increased hepatic fatty acid abundance, consistent with impaired fatty acid utilization and/or enhanced retention within the liver, while the *Tmem135* mutation strongly reduces DHA levels independent of changes in total circulating fatty acids.

**Figure 2.**
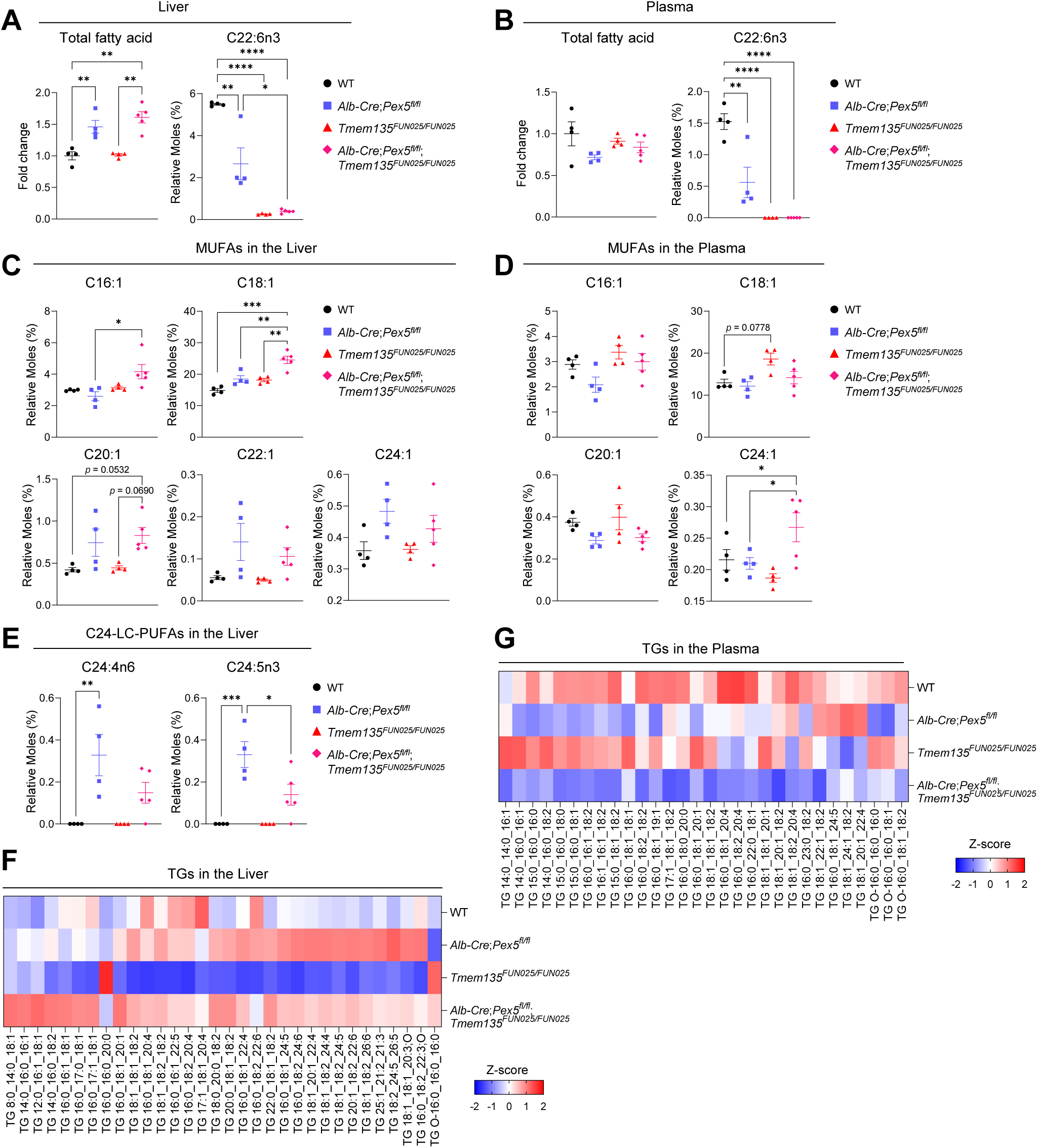
TMEM135 mutation enhances MUFA-rich lipid export, whereas peroxisomal dysfunction promotes hepatic retention. (A) Quantification of total hepatic fatty acid content and docosahexaenoic acid (DHA; C22:6) levels in WT, *Alb-Cre*;*Pex5^fl/fl^*, *Tmem135^FUN025/FUN025^*, and *Alb-Cre*;*Pex5^fl/fl^*;*Tmem135^FUN025/FUN025^* mice. (B) Quantification of total plasma fatty acid content and DHA (C22:6) levels in WT, *Alb-Cre*;*Pex5^fl/fl^*, *Tmem135^FUN025/FUN025^*, and *Alb-Cre*;*Pex5^fl/fl^*;*Tmem135^FUN025/FUN025^* mice. (C) Quantification of hepatic monounsaturated fatty acid (MUFA) levels, including C16:1, C18:1, C20:1, C22:1, and C24:1, in WT, *Alb-Cre*;*Pex5^fl/fl^*, *Tmem135^FUN025/FUN025^*, and *Alb-Cre*;*Pex5^fl/fl^*;*Tmem135^FUN025/FUN025^* mice. (D) Quantification of plasma MUFA levels, including C16:1, C18:1, C20:1, and C24:1, in WT, *Alb-Cre*;*Pex5^fl/fl^*, *Tmem135^FUN025/FUN025^*, and *Alb-Cre*;*Pex5^fl/fl^*;*Tmem135^FUN025/FUN025^* mice. (E) Quantification of hepatic C24-long-chain fatty acid (C24-LC-PUFA) species, including C24:4n6 and C24:5n3, in WT, *Alb-Cre*;*Pex5^fl/fl^*, *Tmem135^FUN025/FUN025^*, and *Alb-Cre*;*Pex5^fl/fl^*;*Tmem135^FUN025/FUN025^* mice. (F) Heatmap showing Z-score-normalized abundance of triglyceride (TG) species in livers from WT, *Alb-Cre*;*Pex5^fl/fl^*, *Tmem135^FUN025/FUN025^*, and *Alb-Cre*;*Pex5^fl/fl^*;*Tmem135^FUN025/FUN025^* mice. (G) Heatmap showing Z-score-normalized abundance of TG species in plasma from WT, *Alb-Cre*;*Pex5^fl/fl^*, *Tmem135^FUN025/FUN025^*, and *Alb-Cre*;*Pex5^fl/fl^*;*Tmem135^FUN025/FUN025^* mice. Fatty acid quantification data are presented as mean ± SEM (n = 4–5 mice per group). Statistical significance was determined by two-way ANOVA followed by Šídák’s multiple-comparisons test. \**p* < 0.05, \*\**p* < 0.01, \*\*\**p* < 0.001, \*\*\*\**p* < 0.0001

Analysis of fatty acid composition revealed a trend toward increased MUFAs, particularly 18:1, in both *Tmem135* mutant and *Pex5*-deficient livers; however, these changes did not reach statistical significance individually. In contrast, 18:1 levels were significantly elevated in the double mutant, indicating a combined effect on MUFA accumulation (Figure 2C and S2A). Similarly, elongation products of 18:1, including 20:1, 22:1, and 24:1, showed increasing trends—most prominently in the double mutant—although these changes were not consistently statistically significant. In plasma, elevated 18:1 levels were observed primarily in the *Tmem135* mutant, but not in *Pex5*-deficient mice (Figure 2D and S2B), suggesting that the *Tmem135* mutation is associated with increased circulating MUFAs, whereas loss of peroxisomal function shifts MUFA distribution toward the liver. In *Pex5*-deficient liver, we observed accumulation of specific C24-long-chain polyunsaturated fatty acids (C24-LC-PUFA), including 24:4n-6 and 24:5n-3 (Figure 2E). Notably, these species are preferentially retained within the liver under conditions of impaired peroxisomal function. Desaturation of 24:5n-3 in the endoplasmic reticulum yields 24:6n-3, which is then transported to peroxisomes, where β-oxidation produces DHA. The accumulation of 24:5n-3 and the loss of DHA in the in *Tmem135* mutant and double-mutant livers support a critical role for peroxisomes in liver DHA biosynthesis. Consistent with changes in fatty acid composition, triglyceride (TG) analysis revealed increased abundance of TG species in *Pex5*-deficient and double-mutant livers (Figure. 2F). In contrast, the *Tmem135* mutant liver exhibited an opposite trend, with many of these TG species showing reduced relative abundance, indicating that elevated MUFA levels in this context are not preferentially directed toward hepatic storage.

In plasma, TG species—particularly those containing 18:1—were enriched in *Tmem135* mutant mice, but not in *Pex5*-deficient animals (Figure 2G), including abundant species such as TG 16:0_18:1_18:1, TG 16:0_18:1_20:1, and TG 18:1_18:1_20:1 (Figure 2G and S3A), consistent with increased incorporation of MUFAs into circulating TGs. In contrast, these MUFA-containing TG species accumulated in the livers of *Pex5*-deficient and double-mutant mice, indicating that loss of peroxisomal function selectively shifts MUFA-containing TGs from circulating plasma pools toward hepatic retention. Notably, hepatic SCD1 expression remained similarly elevated in *Tmem135* mutant and double-mutant livers (Figure S3B), and there was virtually no change in the levels of MUFA-containing lipid species other than TG across these genotypes (Figure S3C and S3D). In addition, CD36 abundance, which was elevated in *Tmem135* mutant liver, was restored toward control levels in the double mutant (Figure S3E), suggesting altered hepatic fatty acid uptake and retention under conditions of peroxisomal dysfunction. In contrast, plasma apolipoprotein abundance remained largely unchanged across genotypes (Figure S3F).

Together, these findings support a model in which the *Tmem135* mutation drives MUFA production and alters hepatic and circulating lipid profiles, while peroxisomal function regulates the partitioning of these lipids between export and storage. Disruption of peroxisomal capacity shifts this balance toward hepatic retention and storage of MUFA- and C24-LC-PUFA-containing TGs.

### Peroxisomal pathways influence lipid partitioning toward triglyceride storage

To investigate whether changes in peroxisomal capacity contribute to the observed lipid phenotypes, we examined peroxisomal abundance and associated metabolic pathways. Quantification of peroxisome revealed an increase in peroxisomal numbers in *Tmem135* mutant liver (Figure S1C)^8^, suggesting enhanced peroxisomal capacity under conditions of the *Tmem135* mutation. In contrast, peroxisomal structures were reduced or disrupted in *Pex5*-deficient hepatocytes, consistent with impaired peroxisomal function. Accordingly, expression of key peroxisomal β-oxidation enzymes, including ACOX1, DBP, and SCPx, was increased in *Tmem135* mutant liver, indicating activation of peroxisomal fatty acid oxidation pathways (Figure 3B, 3C, and 3D). In contrast, these proteins were significantly decreased in *Pex5*-deficient and double-mutant livers, reflecting a loss of functional peroxisomal capacity.

**Figure 3.**
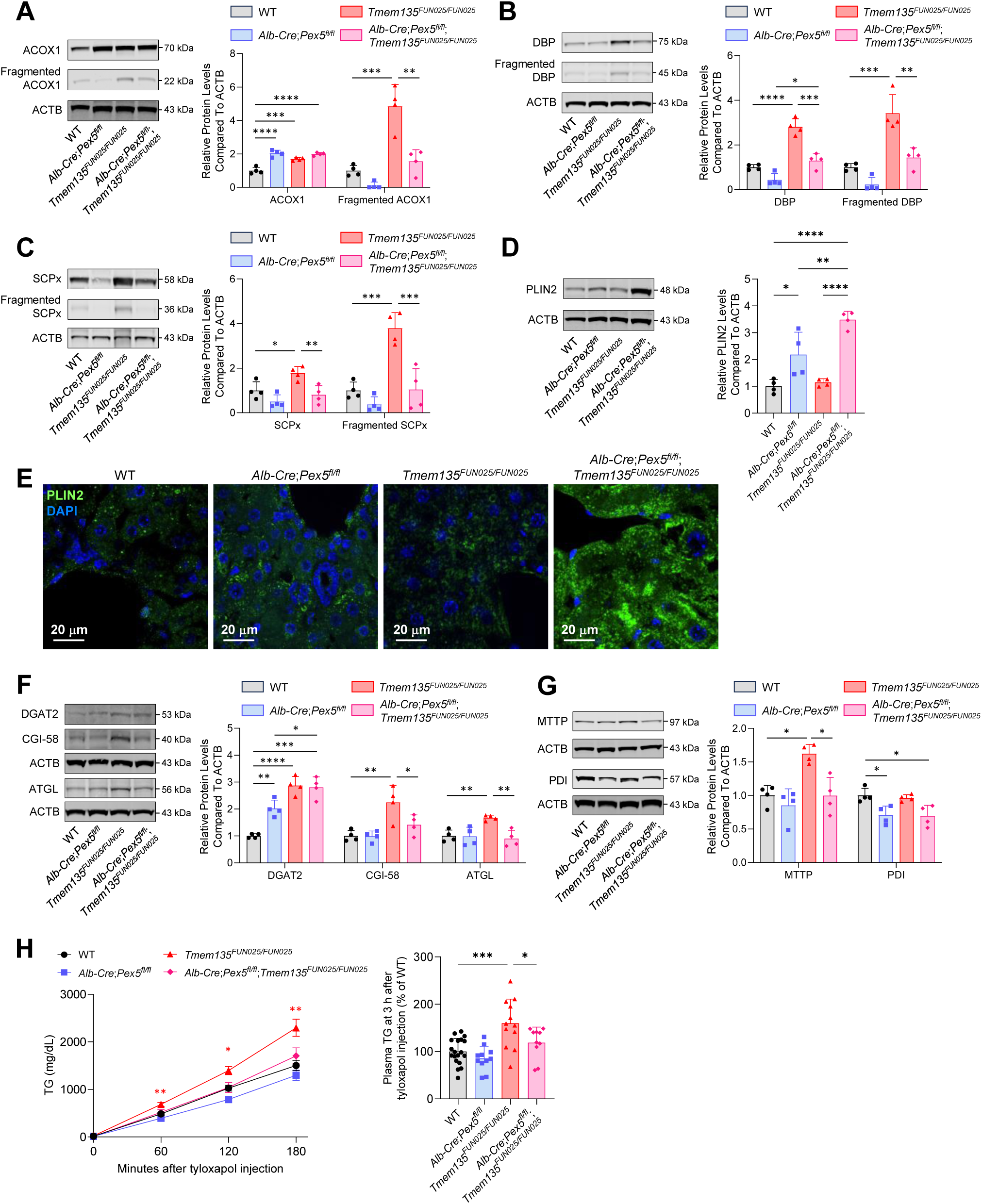
Peroxisomal dysfunction shifts hepatic lipid handling from mobilization to storage. (A) Representative immunoblots and quantification of full-length and fragmented peroxisomal acyl-coenzyme A oxidase 1 (ACOX1) in the liver lysates from 3-month-old WT, *Alb-Cre*;*Pex5^fl/fl^*, *Tmem135^FUN025/FUN025^*, and *Alb-Cre*;*Pex5^fl/fl^*;*Tmem135^FUN025/FUN025^* mice. (B) Representative immunoblots and quantification of full-length and fragmented D-bifunctional protein (DBP) in liver lysates from the indicated genotypes. (C) Representative immunoblots and quantification of full-length and fragmented sterol carrier protein X (SCPx) in liver lysates from the indicated genotypes. (D) Representative immunoblots and quantification of perilipin-2 (PLIN2) in liver lysates from the indicated genotypes. (E) Representative high-resolution confocal micrographs of PLIN2 immunostaining (green) and DAPI (blue) in liver sections from the indicated genotypes. Scale bar, 20 μm. (F) Representative immunoblots and quantification of diacylglycerol *O*-acyltransferase 2 (DGAT2), comparative gene identification-58 (CGI-58), and adipose triglyceride lipase (ATGL) in liver lysates from the indicated genotypes. (G) Representative immunoblots and quantification of microsomal triglyceride transfer protein (MTTP) and protein disulfide isomerase (PDI) in liver lysates from the indicated genotypes. (H) Line graph shows plasma TG levels after tyloxapol injection, presented as mean ± SEM. Red asterisks indicate significant differences between WT and *Tmem135^FUN025/FUN025^* mice. Bar graph shows 3 h endpoint plasma TG levels as mean ± SD (n = 10–19 mice per group). All immunoblot quantification data are presented as mean ± SD (n = 4 mice per group). β-actin (ACTB) served as the loading control for these Western blot experiments. Statistical significance was determined by two-way ANOVA followed by Šídák’s multiple-comparisons test. \**p* < 0.05, \*\**p* < 0.01, \*\*\**p* < 0.001, \*\*\*\**p* < 0.0001

We next examined lipid droplet–associated proteins to assess lipid storage. Expression of PLIN2, a marker of lipid droplet formation and stabilization, was increased in *Pex5*-deficient and double-mutant livers (Figure 3D and 3E), consistent with enhanced lipid storage under conditions of impaired peroxisomal function. In parallel, DGAT2, a rate-limiting enzyme for TG synthesis, was increased in both *Tmem135* mutant and *Pex5*-deficient livers (Figure 3F). In contrast, CGI-58 and ATGL, which promote TG hydrolysis and lipid droplet turnover, were selectively increased in *Tmem135* mutant liver but restored toward control levels in double-mutant mice (Figure 3F), suggesting that loss of peroxisomal function suppresses compensatory lipid catabolic pathways induced by the *Tmem135* mutation. We further examined proteins involved in hepatic lipid export. MTTP, which is required for VLDL assembly and secretion, was increased in *Tmem135* mutant liver but restored toward control levels in the double mutant, while PDI expression was reduced in *Pex5*-deficient liver (Figure 3G). To assess hepatic TG export *in vivo*, we measured plasma TG levels 3 hours after tyloxapol administration, which blocks lipoprotein lipase–mediated TG clearance. Under food-deprived conditions, plasma TG levels were significantly increased in *Tmem135* mutant mice compared with WT mice (Figure 3H), indicating enhanced hepatic VLDL-TG secretion. In contrast, this increase was suppressed in *Alb-Cre*;*Pex5^fl/fl^*;*Tmem135^FUN025/FUN025^* mice. These findings suggest that the TMEM135 mutation promotes hepatic VLDL-TG export, whereas loss of peroxisomal function attenuates this effect, indicating that peroxisomal pathways contribute to TMEM135-dependent regulation of hepatic lipid export.

Together, these results indicate that the *Tmem135* mutation is associated with increased peroxisomal capacity and β-oxidation potential and promotes adaptive pathways involved in TG mobilization and lipid export, whereas loss of PEX5 leads to a reduction in peroxisomal enzyme abundance and function, shifting hepatic lipid handling toward TG retention and storage.

### Proteomic analysis reveals distinct DHA-insensitive and DHA-sensitive metabolic programs in *Tmem135* mutant liver

To gain insight into the metabolic programs underlying the altered lipid profiles and partitioning observed in *Tmem135* mutant liver, we performed quantitative proteomic analysis. In addition to *Tmem135* mutant and control mice, we analyzed *Tmem135* mutant mice supplemented with DHA-rich fish oil (10% w/w) to determine whether restoration of DHA levels alters the associated proteomic profile. Volcano plot analysis revealed widespread changes in protein abundance in *Tmem135* mutant livers compared to WT controls (Figure 4A). In total, 855 proteins were significantly altered, including 513 upregulated and 342 downregulated proteins, indicating broad metabolic remodeling associated with the *Tmem135* mutation.

**Figure 4.**
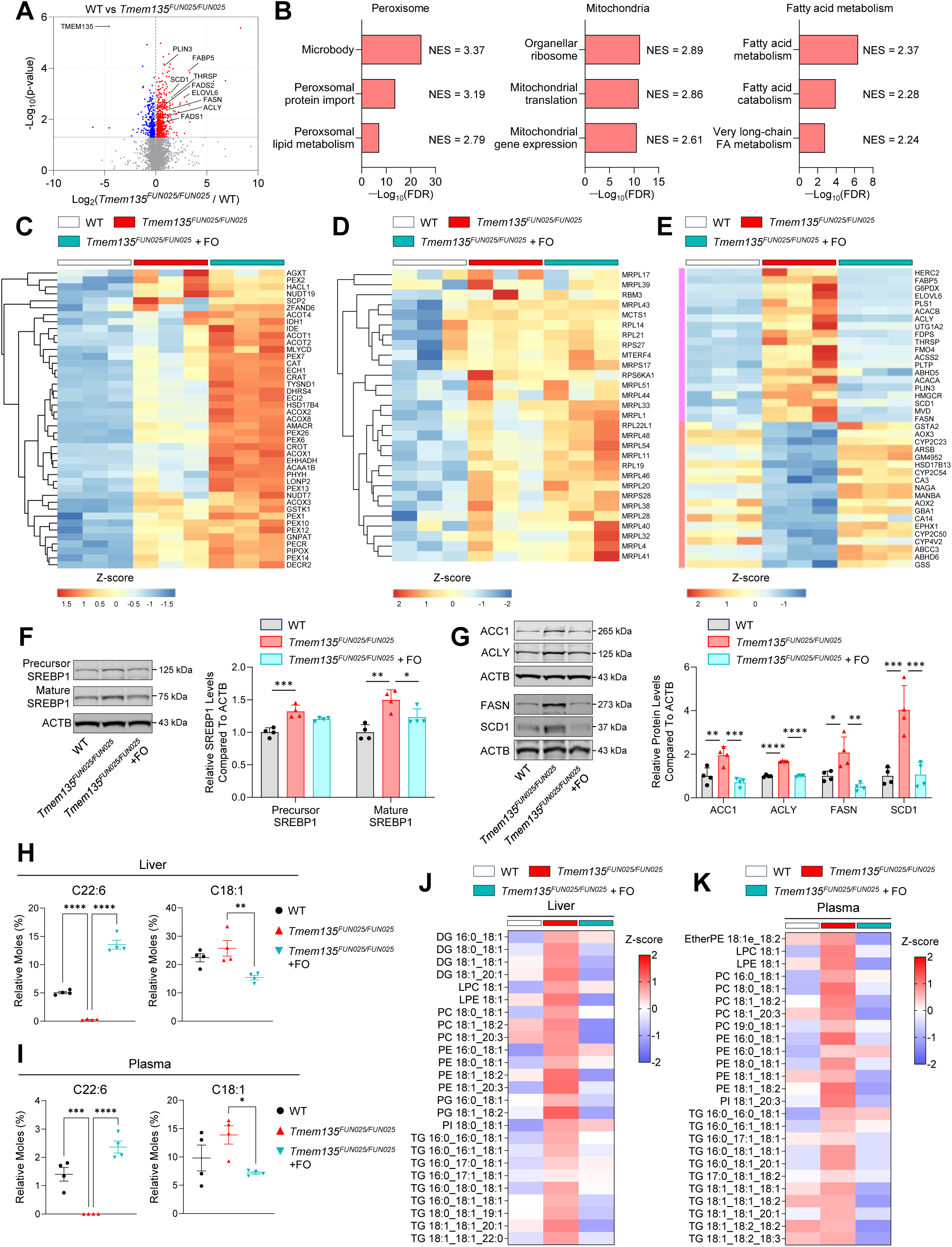
TMEM135 mutation induces distinct DHA-insensitive and DHA-sensitive metabolic programs. (A) Volcano plot showing differentially expressed proteins between 4-month-old WT and *Tmem135^FUN025/FUN025^* livers. Significantly increased and decreased proteins (*p* < 0.05) are highlighted in red and blue, respectively. (B) Gene Set Enrichment Analysis (GSEA) of proteomic data comparing WT and *Tmem135^FUN025/FUN025^* livers. Enriched pathways associated with peroxisomal function, mitochondrial metabolism, and fatty acid metabolism are ranked by false discovery rate (FDR). NES, normalized enrichment score. (C) Heatmap showing Z-score-normalized abundance of peroxisomal proteins identified by proteomic analysis in WT, *Tmem135^FUN025/FUN025^*, and fish oil (FO)-fed *Tmem135^FUN025/FUN025^* livers. (D) Heatmap showing Z-score-normalized abundance of proteins involved in mitochondrial translation in WT, *Tmem135^FUN025/FUN025^*, and FO-fed *Tmem135^FUN025/FUN025^* livers. (E) Heatmap showing Z-score-normalized abundance of proteins altered in *Tmem135^FUN025/FUN025^* livers and restored by FO feeding in the indicated genotypes. Pink and red bars denote proteins increased and decreased, respectively, in *Tmem135^FUN025/FUN025^* livers and restored toward WT levels by FO feeding. (F) Representative immunoblots and quantification of precursor and mature sterol regulatory element-binding protein 1 (SREBP1) in liver lysates from WT, *Tmem135^FUN025/FUN025^*, and FO-fed *Tmem135^FUN025/FUN025^* mice. (G) Representative immunoblots and quantification of the lipogenic enzymes acetyl-CoA carboxylase 1 (ACC1), ATP-citrate lyase (ACYL), fatty acid synthase (FASN), and stearoyl-CoA desaturase 1 (SCD1) in liver lysates from WT, *Tmem135^FUN025/FUN025^*, and FO-fed Tmem135*^FUN025/FUN025^* mice. (H) Quantification of hepatic docosahexaenoic acid (DHA; C22:6) and oleic acid (C18:1) levels in WT, *Tmem135^FUN025/FUN025^*, and FO-fed *Tmem135^FUN025/FUN025^* mice. (I) Quantification of plasma DHA (C22:6) and oleic acid (C18:1) levels in WT, *Tmem135^FUN025/FUN025^*, and FO-fed *Tmem135^FUN025/FUN025^* mice. (J) Heatmap showing Z-score-normalized abundance of C18:1-containing lipid species in livers from WT, *Tmem135^FUN025/FUN025^*, FO-fed *Tmem135^FUN025/FUN025^* mice. (K) Heatmap showing Z-score-normalized abundance of C18:1-containing lipid species in plasma from WT, *Tmem135^FUN025/FUN025^*, FO-fed *Tmem135^FUN025/FUN025^* mice. Immunoblot quantification data are presented as mean ± SD, and β-actin (ACTB) served as the loading control for these Western blot experiments. Fatty acid quantification data are presented as mean ± SEM (n = 4 mice per group). Statistical significance was determined by one-way ANOVA followed by Dunnett’s multiple-comparisons test. \**p* < 0.05, \*\**p* < 0.01, \*\*\**p* < 0.001, \*\*\*\**p* < 0.0001

To investigate the functional relevance of the *Tmem135* mutation, we conducted gene set enrichment analysis (GSEA) to identify pathways differentially enriched between WT and *Tmem135* mutant livers. This analysis revealed significant enrichment of gene sets associated with peroxisomal function, mitochondrial pathways, and fatty acid metabolism in *Tmem135* mutant livers (Figure 4B). Positive normalized enrichment scores (NES) indicated that these pathways were predominantly driven by proteins upregulated in the mutant liver proteome. Heatmap analysis revealed that proteins associated with peroxisomal and mitochondrial metabolism were broadly increased in *Tmem135* mutant liver (Figure 4C, 4D). Immunoblot analysis further validated representative changes in peroxisomal proteins (Figure S4A, and S4B). Notably, these changes were largely insensitive to DHA supplementation, as similar patterns of protein abundance were maintained following DHA-rich fish oil treatment. These results indicate that the *Tmem135* mutation induces a DHA-independent metabolic program characterized by enhanced peroxisomal and mitochondrial metabolic capacity.

In contrast, a distinct subset of proteins displayed marked sensitivity to DHA supplementation (Figure 4E). These proteins segregated into two major functional groups according to their directionality of regulation. First, proteins involved in lipid biosynthesis and SREBP-dependent metabolic pathways, including cholesterol biosynthesis and fatty acid metabolism, were prominently enriched in *Tmem135* mutant liver (Figure 4E and S4C). Consistent with these proteomic findings, the activated cleaved form of SREBP1 was markedly increased in *Tmem135* mutant liver (Figure 4F). Correspondingly, multiple canonical downstream lipogenic enzymes, including ACC1, ACLY, SCD1, and FASN, were elevated in the mutant liver and restored toward WT levels following fish oil supplementation (Figure 4G). Furthermore, proteins involved in lipid droplet turnover and lipid export, including CGI-58, ATGL, and MTTP, were regulated in a DHA-dependent manner (Figure S4D and S4E). These findings indicate that activation of the SREBP1-driven lipogenic program in *Tmem135* mutant liver represents a DHA-dependent metabolic response. A second group of proteins exhibited the opposite DHA-dependent response, being reduced in *Tmem135* mutant liver and restored following fish oil treatment. These proteins were enriched in pathways associated with hepatocellular homeostasis, including xenobiotic metabolism, oxidoreductase activity, and amino acid and glutathione metabolism, consistent with reduced expression of proteins such as CYP2C23, ABCC3, GSTA2, and GSS in the mutant (Figure 4E and S4F). Notably, enrichment of an HNF4A-associated gene set further supports suppression of an HNF4A-linked hepatocellular homeostatic program in the mutant state. Consistent with these proteomic changes, lipidomic analysis demonstrated a significant increase in 18:1 levels in *Tmem135* mutant liver, reflecting enhanced lipogenesis and fatty acid desaturation (Figure 4H, 4I, 4J, 4K, S5, S6A, and S6B). Fish oil supplementation reduced 18:1 levels in both liver and plasma while increasing DHA levels, confirming that DHA availability directly modulates this lipogenic program. In contrast, fish oil supplementation had minimal effects on the abundance of most plasma apolipoproteins, with APOE representing a notable exception, as its levels were restored toward control levels (Figure S6C).

Together, these results identify two distinct but interconnected metabolic programs in *Tmem135* mutant liver: a DHA-insensitive program involving peroxisomal and mitochondrial metabolic pathways, and a DHA-sensitive program that bifurcates into activation of SREBP-dependent lipogenesis and suppression of an HNF4A-associated hepatocellular homeostatic network.

### Global malonylation changes in *Tmem135* mutant liver

Given that increased lipogenesis is associated with elevated malonyl-CoA production^21– 23^, a key substrate for protein malonylation, we next examined whether the *Tmem135* mutation alters the hepatic malonyl-proteome. Immunoblot analysis revealed that the *Tmem135* mutation increased overall protein malonylation levels in the liver (Figure 5A). Notably, DHA-rich fish oil supplementation also increased the total malonylation signal. However, examination of individual bands revealed a heterogeneous pattern, with both increased and decreased signals across different molecular weights, indicating that changes in malonylation are not uniform across proteins but instead reflect selective remodeling of the malonyl-proteome. In contrast, other major post-translational modifications, including acetylation, succinylation, and ubiquitination, were not substantially altered in mutant liver (Figure S7A, S7B, and S7C). Furthermore, the abundance of mitochondrial malonyl-CoA–related proteins, including malonyl-CoA decarboxylase (MCD), responsible for malonyl-CoA degradation, and ACSF3, a mitochondrial malonyl-CoA synthetase, were unchanged in the mutant liver (Figure S7D and S7E). By contrast, sirtuin 5 (SIRT5) expression was increased in *Tmem135* mutant liver and restored toward control levels following fish oil supplementation (Figure S7F), suggesting activation of a compensatory response to altered protein malonylation dynamics.

**Figure 5.**
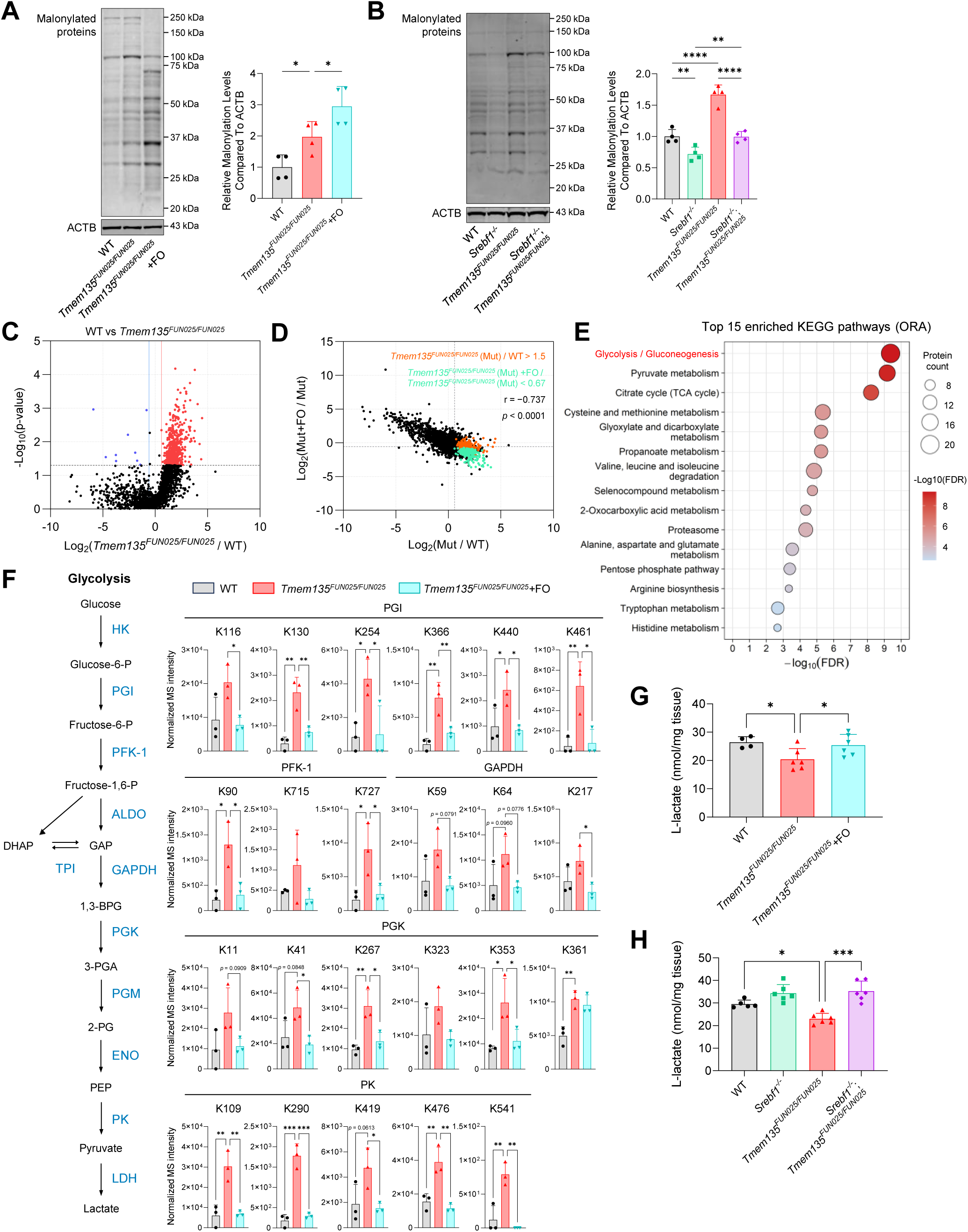
TMEM135 mutation promotes SREBP-dependent protein malonylation and glycolytic remodeling. (A) Representative immunoblots and quantification of malonylated proteins in the liver lysates from WT, *Tmem135^FUN025/FUN025^*, fish oil (FO)-fed *Tmem135^FUN025/FUN025^* mice. (B) Representative immunoblots and quantification of malonylated proteins in the liver lysates from WT, *Srebf1^-/-^*, *Tmem135^FUN025/FUN025^*, *Srebf1^-/-^*;*Tmem135^FUN025/FUN025^* mice. (C) Representative immunoblots and quantification of full-length and fragmented sterol carrier protein X (SCPx) in liver lysates from the indicated genotypes. (D) Scatter plot of malonylation sites comparing changes associated with *Tmem135* mutation and fish oil (FO) feeding. The x axis represents log₂(*Tmem135^FUN025/FUN025^* (Mut)/WT), and the y axis represents log₂(FO-fed *Tmem135^FUN025/FUN025^* (Mut+FO) / *Tmem135^FUN025/FUN025^* (Mut)). Orange dots indicate sites increased in *Tmem135^FUN025/FUN025^* livers relative to WT (fold change > 1.5), whereas green dots indicate sites decreased following FO feeding relative to *Tmem135^FUN025/FUN025^* controls (fold change < 0.67). (E) KEGG pathway over-representation analysis (ORA) of proteins harboring malonylation sites that were increased in *Tmem135^FUN025/FUN025^* livers and restored by FO feeding, independent of changes in total protein abundance. (F) Schematic illustration of the glycolytic pathway and quantification of malonylated lysine sites in phosphoglucose isomerase (PGI), phosphofructokinase-1 (PFK-1), glyceraldehyde-3-phosphate dehydrogenase (GAPDH), phosphoglycerate kinase (PGK), and pyruvate kinase (PK) in WT, *Tmem135^FUN025/FUN025^*, and FO-fed *Tmem135^FUN025/FUN025^* mice. (G) L-lactate levels in the livers of WT, *Tmem135^FUN025/FUN025^*, and FO-fedTmem135*^FUN025/FUN025^* mice. **(H)** L-lactate levels in the livers of WT, *Srebf1^-/-^*, *Tmem135^FUN025/FUN025^*, *Srebf1^-^* ^/-^;Tmem135^FUN025/FUN025^ mice. All bar graphs are presented as mean ± SD (n = 4–6 mice per group). β-actin (ACTB) served as the loading control for Western blot experiments. Statistical significance was determined by one-way ANOVA followed by Dunnett’s multiple-comparisons test for comparisons among WT, *Tmem135^FUN025/FUN025^*, and FO-fed *Tmem135^FUN025/FUN025^* mice, and by two-way ANOVA followed by Šídák’s multiple-comparisons test for comparisons among WT, *Srebf1^-/-^*, *Tmem135^FUN025/FUN025^*, *Srebf1^-/-^*;*Tmem135^FUN025/FUN025^* mice. \**p* < 0.05, \*\**p* < 0.01, \*\*\**p* < 0.001, \*\*\*\**p* < 0.0001

To determine whether these changes are linked to activation of the lipogenic program, we next examined malonylation in *Tmem135* mutant liver in the context of SREBP inhibition. Suppression of SREBP activity reduced activation of canonical SREBP1 downstream pathways (Figure S8A), accompanied by attenuation of the elevated protein malonylation observed in *Tmem135* mutant liver (Figure 5B). These findings indicate that TMEM135-dependent remodeling of the hepatic malonyl-proteome is closely associated with activation of the SREBP-driven lipogenic program.

To define these changes at site-level resolution, we performed quantitative malonyl-proteomic analysis. We identified 4,602 malonylation sites across 1,341 proteins in WT, *Tmem135* mutant, and fish oil (FO) treatment groups (Figure S8B and S8C). Among these, 3,080 sites corresponding to 1,112 proteins were commonly detected in all three groups. Quantitative malonyl-proteomic analysis identified 547 significantly altered malonylation sites between WT and *Tmem135* mutant liver (criteria: [fold-change > 1.5 or < 0.67, p < 0.05]), including 538 increased and 9 decreased sites, revealing a strong bias toward increased malonylation in the mutant (Figure 5C). Scatter plot analysis of the 3,080 commonly detected sites, comparing log2(*Tmem135^FUN025/FUN025^* (Mut) / WT) with log2(Mut+FO/Mut), revealed a strong inverse correlation (r = −0.727), indicating that mutant-induced malonylation remodeling is broadly opposed by DHA supplementation (Figure 5D). Stringent filtering further identified a defined subset of DHA-sensitive malonylation events that were increased in *Tmem135* mutant liver and restored following FO supplementation (Mut/WT > 1.5-fold, Mut+FO/Mut < 0.67, both p < 0.05). In contrast, although mitochondrial proteins are known to undergo extensive malonylation^24^, mitochondrial protein malonylation was largely insensitive to FO supplementation, suggesting that DHA-responsive remodeling selectively affects specific subsets of the hepatic malonyl-proteome (Figure S8D and S8E).

To define biological pathways selectively associated with mutant-driven malonylation remodeling, we next focused on DHA-sensitive malonylation events identified by stringent filtering and further extracted sites whose malonylation changes were not correlated with alterations in protein abundance (protein abundance correlation FDR > 0.20). KEGG pathway analysis of these protein abundance–independent malonylation events revealed prominent enrichment of glycolytic pathways, indicating that glycolytic enzymes undergo coordinated hyper-malonylation in *Tmem135* mutant liver independently of altered protein expression (Figure 5E, 5F, S8F and S8G). To assess the functional consequences of glycolytic enzyme hypermalonylation, we measured hepatic L-lactate levels. *Tmem135* mutant liver exhibited a significant reduction in L-lactate levels, whereas FO supplementation restored lactate levels toward those of WT controls (Figure 5G). Notably, *Srebf1* deficiency similarly rescued the reduction in L-lactate levels observed in *Tmem135* mutant liver (Figure 5H).

Taken together, these data support a model in which the *Tmem135* mutation reduces DHA availability, leading to activation of a lipogenic program and increased malonyl-CoA production. Elevated malonyl-CoA is associated with selective protein malonylation, which modulates glycolytic pathways at the post-translational level. In this framework, the *Tmem135* mutation establishes a lipogenic metabolic state in which peroxisomal function determines whether lipid homeostasis is maintained or shifted toward storage. Concurrently, increased malonylation provides a mechanism to coordinate broader metabolic adaptation across cellular pathways (Figure 6).

**Figure 6.**
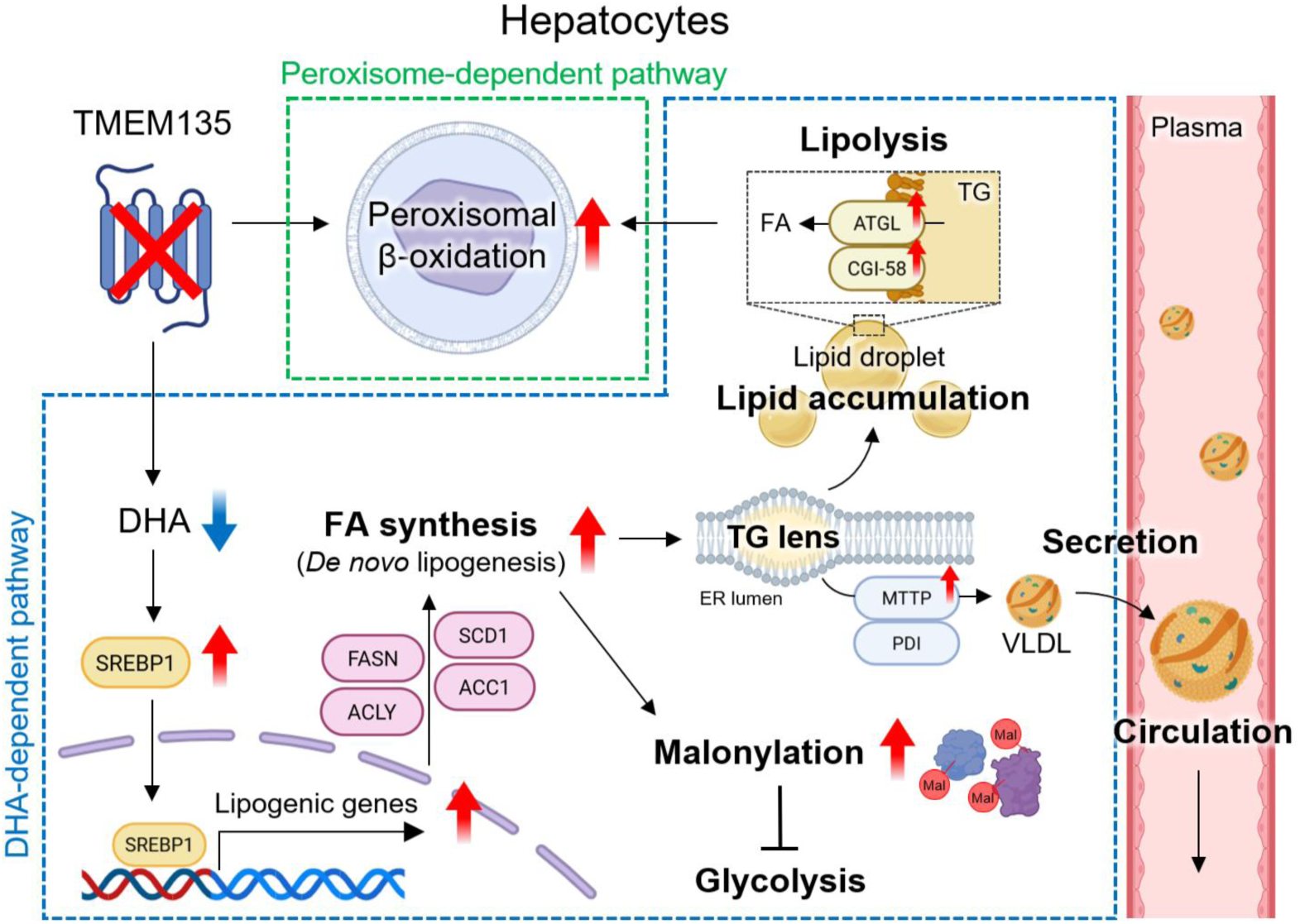
TMEM135 deficiency remodels hepatic lipogenesis and protein malonylation through a DHA-sensitive program buffered by peroxisomal function.

## Discussion

In this study, we demonstrate that the *Tmem135* mutation establishes a lipogenic hepatic state that is buffered by peroxisomal metabolic capacity. Using a genetic interaction approach combining the *Tmem135* mutation with liver-specific *Pex5* deletion, we show that disruption of peroxisomal function reveals a latent metabolic imbalance, resulting in hepatic lipid accumulation. Integrated lipidomic and proteomic analyses further reveal that this metabolic state consists of distinct DHA-insensitive and DHA-sensitive programs, including activation of SREBP-dependent lipogenesis and suppression of a hepatocellular homeostatic network.

The genetic interaction between the *Tmem135* mutation and *Pex5* deficiency provides key insight into the role of peroxisomal function in regulating hepatic lipid homeostasis. While the *Tmem135* mutation alone promotes a lipogenic state, it does not result in hepatic steatosis, indicating the presence of compensatory mechanisms. In contrast, loss of PEX5-dependent peroxisomal protein import in the *Tmem135* mutant background leads to marked lipid accumulation, demonstrating that peroxisomal metabolic capacity buffers elevated lipid storage. Previous work in our model has shown that the *Tmem135* mutation increases peroxisome abundance through PPARα signaling^8^. In this study, we further show that this pathway operates independently of DHA availability. Supporting this, the increase in peroxisomal and mitochondrial proteins observed in *Tmem135* mutant liver was largely insensitive to DHA-rich fish oil supplementation, suggesting that organelle-level metabolic capacity represents a distinct, DHA-independent buffering mechanism. Given that TMEM135 has also been implicated in mitochondrial dynamics^6,25^, and consistent with this, we observed elongated mitochondrial morphology in *Tmem135* mutant liver^5^, which suggests that coordinated changes in peroxisomal and mitochondrial function may further contribute to maintaining metabolic balance under conditions of increased lipogenesis.

Our findings further suggest that peroxisomal function plays a critical role in determining lipid partitioning under conditions of increased lipogenesis. Notably, disruption of PEX5 preferentially promoted accumulation of TGs enriched in MUFAs, particularly 18:1, indicating that lipid allocation is not uniform across fatty acid species. Among the lipid species analyzed, 18:1 most closely tracked the phenotypic pattern of the genetic interaction, suggesting that MUFA metabolism represents a key node in this process, probably due to the increased expression of the enzymes involved in MUFA biosynthesis. This specificity may reflect the central role of FASN-mediated biosynthesis of 16:0, which is elongated to 18:0 for subsequent SCD1-mediated desaturation to generate 18:1^26,27^, which serves as a preferred substrate for TG synthesis, or differential routing of MUFAs into storage versus membrane lipid pools. Consistent with this, increased expression of lipid droplet–associated proteins such as PLIN2^28^ in *Pex5*-deficient and double-mutant liver may facilitate sequestration of MUFA-rich lipids into TGs. Conversely, increased MTTP expression and tyloxapol-based TG secretion analysis suggest that the *Tmem135* mutation enhances VLDL-TG secretion into circulation, whereas loss of peroxisomal function attenuates this export pathway.

The *Tmem135* mutation induces broad remodeling of hepatic lipid metabolism, characterized by activation of SREBP-dependent lipogenesis. DHA is a well-established regulator of SREBP1 activity^9–11^, and our findings recapitulate this regulatory axis *in vivo*. In *Tmem135* mutant liver, increased expression of lipogenic enzymes, including SCD1, is associated with elevated levels of 18:1, indicating enhanced fatty acid synthesis and desaturation. Fish oil supplementation suppresses this program, confirming that DHA availability directly regulates this lipogenic state. Although fish oil contains both DHA and EPA, our findings suggest that the observed effects are more closely associated with restoration of DHA than with changes in EPA abundance. Hepatic EPA levels were largely unchanged in *Tmem135* mutant and *Pex5*-deficient *Tmem135* mutant mice, whereas DHA was consistently reduced and restored by fish oil supplementation. These observations are consistent with a DHA-sensitive mechanism linking TMEM135 deficiency to activation of SREBP-dependent lipogenesis. These results support a model in which reduced DHA availability in *Tmem135* mutant liver drives activation of SREBP-dependent lipogenesis, leading to increased MUFA production.

In parallel with activation of lipogenesis, we identified suppression of a distinct set of pathways associated with hepatocellular homeostasis. These include pathways involved in xenobiotic metabolism^29^, redox balance^30,31^, and amino acid metabolism^32^, which were reduced in the *Tmem135* mutant and restored by DHA supplementation. Notably, this group includes HSD17B13, a lipid droplet–associated protein^33^ genetically linked to susceptibility to NAFLD^34^, suggesting potential clinical relevance of this regulatory axis. Enrichment of an HNF4A-associated gene set^35^ suggests that the *Tmem135* mutation suppresses a broader hepatocellular homeostatic program, although HNF4A abundance itself was not altered. This finding indicates that TMEM135-dependent metabolic remodeling may influence hepatocyte functional output through mechanisms that extend beyond direct changes in transcription factor expression.

The *Tmem135* mutant mouse represents a unique model for studying the role of DHA in metabolic regulation *in vivo*. Unlike models of global peroxisomal dysfunction, such as PEX5 deficiency, which broadly disrupt multiple lipid metabolic pathways, the *Tmem135* mutation primarily impairs DHA availability, providing a more defined system to assess the specific contribution of DHA to hepatic metabolic regulation. In this model, DHA levels are markedly reduced in both liver and plasma, allowing direct assessment of the consequences of chronic DHA deficiency. Our findings, together with DHA supplementation experiments, further support a critical role for TMEM135 in endogenous DHA production^8^ and demonstrate that restoration of DHA levels is sufficient to suppress lipogenic pathways and restore broader metabolic homeostasis. These observations highlight the importance of DHA availability in maintaining hepatic metabolic balance. Consistent with this, clinical and preclinical studies have shown that DHA supplementation can reduce hepatic lipid accumulation^36–38^ and improve features of NAFLD^39–41^. Our findings provide mechanistic insight into these effects, suggesting that DHA regulates lipid distribution by suppressing lipogenic state and promoting hepatocellular metabolic homeostasis.

In addition to transcriptional and metabolic changes, we identified increased protein malonylation as a downstream consequence of TMEM135-dependent lipogenesis. Protein malonylation is a post-translational modification linked to cellular malonyl-CoA levels^42^, which are elevated during active fatty acid synthesis^21^. Notably, a subset of malonylation changes occurred independently of changes in protein abundance and were sensitive to DHA supplementation, suggesting that lipogenesis introduces an additional layer of metabolic regulation at the post-translational level. Among the proteins exhibiting DHA-sensitive malonylation, glycolytic enzymes were particularly enriched. Previous studies have similarly reported extensive malonylation of glycolytic proteins in SIRT5-deficient, *db*/*db*, and *ob*/*ob* mice^24,43,44^, suggesting that glycolysis is especially susceptible to regulation by malonylation. We also identified DHA-sensitive malonylation of proteins involved in amino acid metabolism, another pathway reported to be highly malonylated in *db*/*db* mice^43^. The convergence of these findings suggests that glycolytic and amino acid metabolic pathways may represent preferential targets of malonylation during conditions of increased lipogenesis. Notably, DHA supplementation has been shown to alleviate inflammation and insulin resistance in *db*/*db* mice^45^, raising the possibility that reversal of DHA-sensitive protein malonylation contributes to the beneficial metabolic effects of DHA. In contrast, although many mitochondrial proteins were also malonylated, these modifications were largely insensitive to DHA supplementation, indicating that malonylation is not uniformly regulated across the proteome. Together, our findings suggest that selective remodeling of malonylation-sensitive metabolic networks may represent a conserved mechanism linking excessive lipogenesis to metabolic dysfunction.

This study has several limitations. While our analyses identify coordinated changes in metabolic pathways and support a model in which TMEM135 regulates DHA availability and lipid metabolism, the mechanisms linking these changes to downstream regulatory programs remain to be fully defined. In particular, how TMEM135-dependent remodeling influences the HNF4A-associated hepatocellular homeostatic program remains unclear. In addition, although increased protein malonylation was observed, the functional consequences of these modifications, including their impact on enzymatic activity and metabolic regulation, require further investigation. Finally, the signaling pathways that connect peroxisomal metabolic capacity to lipid partitioning and hepatocellular homeostasis remain to be elucidated.

In summary, our findings establish TMEM135 as a central regulator of hepatic lipid metabolism and identify peroxisomal function as a key buffering system that determines whether increased lipogenesis is accommodated or redirected toward lipid storage. These results reveal a coordinated metabolic framework in which DHA availability, lipogenesis, and post-translational modification interact to regulate hepatic metabolic state, providing new insight into mechanisms that maintain lipid homeostasis under conditions of metabolic stress.

## DATA AVAILABILITY

Lipidomics data have been deposited in Dryad (accession code: doi:10.5061/dryad.bzkh189s7), and proteomics data have been deposited the ProteomeXchange Consortium via the PRIDE partner repository with the dataset identifier PXD080132.

## ACKNOWLEDGEMENTS

The authors thank Toshi Kinoshita and the University of Wisconsin (UW) Translational Research Initiatives in Pathology laboratory (TRIP), supported by the UW Department of Pathology and Laboratory Medicine, UWCCC (P30 CA014520), and the Office of the Director-NIH (S10OD023526) for access to facilities and services. The authors would like to thank the Biochemistry Optical Core in the Department of Biochemistry at the University of Wisconsin– Madison for imaging support using the SoRa/W1 Spinning Disk Microscope. We are grateful to the UW Biotechnology Center’s Advanced Lipidomics Platform for their time and effort in optimizing the protocols for the lipidomics experiments. The authors would also like to thank Dr. Guanghui Han and PTM BIO for their support with proteomics sample processing and data acquisition, and Research to Prevent Blindness Unrestricted Funds and P30 EY021725 to the Dean McGee Eye Institute, which supported the fatty acid analyses work in the Agbaga Lab. This work was supported by the Timothy William Trout Chair in Eye Research, McPherson Eye Research Institute (to A. Ikeda) and grants from the National Eye Institute (R01 EY030513 to M.P. Agbaga, R01 EY022086 to A. Ikeda, P30 EY016665 to the Department of Ophthalmology and Visual Sciences at the University of Wisconsin-Madison)

## AUTHOR CONTRIBUTIONS

Conceptualization – R.H., M.L., and A.I.; Data curation – R.H., M.L., S.S., P.G., R.S.B., S.M.B., and G.B.W.; Formal analysis – R.H., M.L., P.G., R.S.B., and S.M.B.; Funding Acquisition – M.P.A., and A.I.; Investigation – R.H., M.L., S.S., P.G., and G.B.W.; Methodology –R.H., M.L., S.S., P.G., R.S.B., S.M.B., G.B.W., S.I., C.L.E.Y., T.T., M.P.A., and A.I.; Project Administration – R.H., M.L., S.I., and A.I.; Resources – S.I., A.I.; Supervision – S.I., K.Y., C.L.E.Y., M.P.A., and A.I.; Validation – R.H., M.L., S.S., P.G., R.S.B., S.M.B., G.B.W., S.I., K.Y., C.L.E.Y., T.T., M.P.A., and A.I.; Visualization – R.H., M.L., S.S., and P.G.; Writing-Original Draft – R.H., S.S, S.I., and A.I.; Writing-Review and Editing – R.H., M.L., S.S., P.G., R.S.B., S.M.B., G.B.W., S.I., K.Y., C.L.E.Y., T.T., M.P.A., and A.I.

## DECLARATION OF INTERESTS

The authors declare no competing interests.

## EXPERIMENTAL MODEL AND STUDY PARTICIPANT DETAILS

### Animal models

*Tmem135^FUN025/FUN025^* mice were generated by the previous methods^25^. Albumin-Cre (B6.Cg-*Speer6-ps1^Tg(Alb-cre)21Mgn^*/J), B6J.Pex5-loxP (B6J.129-*Pex5^tm1Pec^*/BaesJ), and SREBP-1c-(B6;129S6-*Srebf1^tm1Mbr^*/J) were obtained from the Jackson Laboratory. These strains were intercrossed to generate *Alb-cre*;*Pex5^fl/fl^*;*Tmem135^FUN025/FUN025^* and *Srebf1^-/-^*;*Tmem135^FUN025/FUN025^* mice on the C57BL/6J background for use in this study. Age-matched wild-type C57BL/6J mice were used as controls. Animals were maintained in the same animal facility at the University of Wisconsin–Madison under standardized housing conditions with a 12-h light/12-h dark cycle (lights on at 6:00 a.m. and off at 6:00 p.m.). All mice were fed a Tekland global soy protein-free extruded rodent diet (#2020X; Inotiv) unless otherwise indicated. For fish oil supplementation, mice a custom fish oil-supplemented diet containing 10% fish oil (w/w) (#TD.230589; Inotiv) from weaning until tissue collection. Both male and female mice at 3 and 4 months of age were included in the study. Unless otherwise indicated, tissue collection and associated experimental procedures were performed during the light phase within a consistent midday interval (approximately 11:00 a.m. to 1:00 p.m.) to reduce potential circadian effects. All animal procedures were carried out in accordance with the National Institutes of Health Guide for the Care and Use of Laboratory Animals and were approved by the Animal Care and Use Committee at the University of Wisconsin–Madison. Experimental procedures and reporting complied with ARRIVE guidelines.

## METHOD DETAILS

### H&E staining

Mice were perfused with a fixative containing 2% glutaraldehyde and 2% paraformaldehyde before the collection of the largest liver lobe for paraffin embedding. Tissue processing and sectioning were performed by the Translational Research Initiatives in Pathology (TRIP) core at the University of Wisconsin–Madison. Paraffin sections were stained with hematoxylin and eosin (H&E) according to the standard protocol.

### Immunohistochemistry

Tissues were fixed overnight at 4 °C in 4% paraformaldehyde, cryoprotected, and embedded in Tissue-Tek O.C.T. Compound. Cryosections were cut at a thickness of 8 µm using a cryostat. Sections were stained with antibodies against PMP70 (#ab3421, Abcam, 1:200 dilution), PEX14 (#10594-1-AP, Proteintech, 1:200 dilution), and PLIN2 to visualize peroxisomes and lipid droplet–associated structures. Secondary donkey anti-rabbit IgG antibodies conjugated to Alexa Fluor 488 (#A21206, Invitrogen) were applied for 45 min at room temperature. Neutral lipids were labeled using BODIPY (#25892, Cayman Cemical) or LipidTOX reagents (#H34476, Invitrogen), and nuclei were counterstained with 4′,6-Diamidine-2′-phenylindole dihydrochloride (DAPI) (#D9542, Sigma Aldrich, 1:1000 dilution). Images of PMP70, PEX14, and BODIPY staining were acquired using a Nikon A1RS confocal microscope at the University of Wisconsin–Madison Optical Imaging Core. PLIN2 and LipidTOX images were acquired using SoRa/W1 Spinning Disk Microscope (Nikon, Tokyo, Japan) at the University of Wisconsin–Madison Biochemistry Optical Core.

### Hepatic TG assay

Liver samples were collected, snap-frozen, and stored at −80 °C until lipid extraction. Briefly, liver tissues were homogenized in Folch (CHCl_3_:MeOH, 2:1), and lipid extracts were dried under nitrogen, resuspended in NP40 substitute assay reagent. TG concentrations were determined using the FUJIFILM Wako L-Type Triglyceride M enzymatic assay (#994-02891), and is normalized by liver weight.

### GC–FID-based fatty acid profiling

Total lipids from liver tissues were extracted using the Folch method^46^, whereas lipids from plasma samples (100 µL) were isolated according to the modified Bligh and Dyer method ^47,48^. Lipid extracts were supplemented with 25 nmol each of 15:0 and 17:0 fatty acids as internal standards. Fatty acid methyl esters (FAMEs) were generated by acid-catalyzed hydrolysis/methanolysis in 16% (v/v) HCl-methanol at 100 °C for 2 h. FAMEs were subsequently extracted with hexane, purified by thin-layer chromatography (TLC), and quantified using an Agilent 7890B gas chromatograph equipped with a flame ionization detector. Fatty acid composition was expressed as relative mol% of total fatty acids.

### Sample preparation for lipidomic analysis

Liver tissues and plasma samples were collected and stored at −80 °C until lipid extraction. All solvents and reagents were pre-chilled on ice prior to use. Liver tissues were handled and sectioned on a dry ice–cooled stainless-steel surface and weighed to the nearest 0.01 mg using a Mettler XSR205 analytical balance. Tissue weights were used for downstream normalization. For plasma lipidomics, 50 µL aliquots from each mouse were analyzed without additional normalization, as equal sample volumes were used throughout the study. Liver tissues were transferred into Qiagen PowerBead tubes (P/N 13113-50) for homogenization and extraction. Lipids were extracted using a solvent mixture consisting of 250 µL PBS, 225 µL methanol containing internal standards (10 µL Avanti SPLASH LipidoMix, Lot#3307-07), and 750 µL methyl tert-butyl ether (MTBE). Samples were homogenized using a Qiagen TissueLyzer II at 30 Hz for three 30-s cycles interspersed with 5-min cooling periods on ice, followed by a final 15-min incubation on ice. Following centrifugation at 16,000 × g for 5 min at 4 °C, 500 µL of the upper organic phase was collected and evaporated to dryness using a Savant SpeedVac concentrator. Lipid extracts were subsequently resuspended in 150 µL isopropanol prior to analysis. A process blank was prepared in parallel with all samples and analyzed concurrently to monitor background contamination. In addition, pooled quality control samples were generated by combining equal volumes from each final lipid extract following resuspension in isopropanol.

### LC–MS/MS-based lipidomic profiling

Samples were analyzed at tissue- and polarity-specific dilution factors. Prior to LC–MS analysis, lipid extracts were diluted in isopropanol by combining the appropriate sample volume with isopropanol in LC–MS vials containing deactivated glass inserts (Agilent P/N 5182-0554 and 5183-2086). Samples were vortexed, briefly centrifuged to collect the solution, and placed in the autosampler for analysis. Lipid separation was performed using a Waters Acquity UPLC BEH C18 column (1.7 µm, 2.1 × 100 mm) coupled to a Waters Acquity UPLC BEH C18 VanGuard pre-column (1.7 µm, 2.1 × 5 mm) maintained at 50 °C. The LC system consisted of an Agilent HiP 1290 Multisampler, Agilent 1290 Infinity II binary pump, and column compartment interfaced with an Agilent 6546 Accurate Mass Q-TOF mass spectrometer equipped with a dual electrospray ionization (ESI) source. For positive ion mode acquisition, source parameters were set as follows: gas temperature 250 °C, gas flow 12 L/min, nebulizer pressure 35 psig, capillary voltage 4000 V, fragmentor voltage 145 V, skimmer voltage 45 V, and octopole RF peak 750 V. For negative ion mode, the gas temperature was set to 350 °C with a drying gas flow of 12 L/min, nebulizer pressure of 25 psig, capillary voltage of 5000 V, fragmentor voltage of 200 V, skimmer voltage of 45 V, and octopole RF peak of 750 V. Reference masses were continuously infused through the second sprayer of the dual ESI source at 15 µL/min using an isocratic pump. Positive ion reference masses were m/z 121.0509 and 922.0098, whereas negative ion reference masses were m/z 112.9856 and 966.0007. Samples were analyzed in randomized order in separate positive and negative ionization experiments over a scan range of m/z 100–1500. Mobile phase A consisted of acetonitrile/water (60:40, v/v) containing 10 mM ammonium formate and 0.1% formic acid, whereas mobile phase B consisted of isopropanol/acetonitrile/water (90:9:1, v/v) containing 10 mM ammonium formate and 0.1% formic acid. The chromatographic gradient began at 15% B, increased to 30% B over 2.4 min, then to 48% B from 2.4–3.0 min, followed by an increase to 82% B from 3.0–13.2 min, and finally to 99% B from 13.2–13.8 min. The gradient was held at 99% B until 15.4 min before returning to initial conditions and equilibrating for 4 min. The flow rate was maintained at 0.5 mL/min throughout the run. Injection volumes for MS1 acquisition were 2 µL in positive ion mode and 5 µL in negative ion mode, whereas MS/MS acquisition volumes were 4 µL and 7 µL for positive and negative ion modes, respectively. Tandem mass spectrometry was performed using the same LC gradient with a narrow isolation width (∼1.3 m/z) and a collision energy of 25 V. MS/MS data acquisition was performed on pooled tissue samples using an iterative acquisition strategy in which each pooled sample was analyzed five times with distinct precursor selections per injection, thereby improving coverage of low-abundance lipid species in the presence of highly abundant lipids.

### Lipidomic data analysis

Lipid species were identified from pooled MS/MS datasets using Lipid Annotator (Agilent) based on accurate precursor masses and diagnostic fragment ions. Identifications were generated independently for positive- and negative-ion modes and included either acyl-chain composition or, where structural resolution was not possible, total carbon number and unsaturation. Quantitative analysis was performed on individual sample MS datasets using MassHunter Profinder (Agilent), which extracted and integrated ion chromatograms based on matched retention time and accurate mass information. Peak area matrices were exported to MetaboAnalyst 5.0 for statistical testing^49^. All lipidomics data have been deposited in the Dryad Digital Repositor (accession code: doi:10.5061/dryad.bzkh189s7).

### Western blotting

Tissues were collected from mice and stored at −80 °C until use. Liver tissues were homogenized in RIPA buffer (#P189901, Thermo Fisher Scientific, Waltham, MA) supplemented with protease inhibitors (#11836170001, Thermo Fisher Scientific) using an Ultra-Turrax T8 tissue homogenizer. Protein concentrations were determined using a BCA Protein Assay Kit (#P123228, Thermo Fisher Scientific). Equal protein amounts were aliquoted, reduced with XT Reducing Agent (#1610792, Bio-Rad, Hercules, CA) for 7 min at 105 °C, separated on 10% Bis-Tris Criterion XT gels (#3450112, Bio-Rad) in MOPS running buffer (#1610788, Bio-Rad) or MES running buffer (#1610789, Bio-Rad), and transferred onto PVDF membranes (#1620177, Bio-Rad) or nitrocellulose membranes (#1620115, Bio-Rad). Membranes were blocked with 5% milk in TBST and incubated overnight at 4 °C with the appropriate primary antibodies at working dilutions of 1:500 or 1:1000. Primary antibodies are listed in supplementary table 1. Following TBST washes, membranes were incubated with the corresponding secondary antibodies at a dilution of 1:5000 (listed in supplementary table 1). Membranes were washed again with TBST and imaged using the Odyssey Imaging System (LI-COR Biosciences, Lincoln, NE). Band intensities were quantified using ImageJ software (NIH, Bethesda, MD). Membranes were stripped using NewBlot Stripping Buffer (LI-COR Biosciences) according to the manufacturer’s instructions and reprobed with additional primary antibodies as required. All immunoband intensities were normalized to the corresponding loading control or to total protein levels determined by Ponceau S staining for each blot.

### VLDL secretion assay

Mice were fasted for 4 hours starting at 7:00 AM. Tyloxapol (#T-0307,Sigma) was dissolved in isotonic saline at a concentration of 20% (v/v) and administered by retro-orbital injection at a dose of 500 mg/kg body weight at 11:00 AM. Blood samples were collected from the tail vein at 0 hour, immediately before tyloxapol administration, and at 1, 2, and 3 hours after administration. Plasma was isolated by centrifugation, and plasma TG concentrations were determined using the FUJIFILM Wako L-Type Triglyceride M enzymatic assay (#994-02891) according to the manufacturer’s instructions. The 3-hour time point was used for endpoint analysis of plasma TG accumulation after tyloxapol administration.

### Protein extraction and enzyme digestion for proteomics

Frozen liver tissues were homogenized and lysed in SDS-containing lysis buffer (8 M urea, 5% SDS, 1× protease inhibitor cocktail, and 50 mM TEAB, pH ∼8) and vortexed thoroughly prior to protein quantification. Protein concentrations were determined using the Pierce™ BCA Protein Assay Kit (Thermo Fisher Scientific). Equal amounts of protein from each sample were subjected to reduction with dithiothreitol (DTT) at 37°C for 60 min, followed by alkylation with iodoacetamide (IAM) at room temperature in the dark for 45 min. Samples were subsequently acidified with phosphoric acid to approximately 2% (pH ≤ 1) according to the manufacturer’s S-Trap™ protocol (Protifi). Acidified lysates were diluted with wash/binding buffer (100 mM TEAB in methanol, pH ∼8), thoroughly mixed, and loaded onto S-Trap™ columns. Columns were washed four times with wash/binding buffer before on-column digestion using protease at a 1:10 enzyme-to-protein ratio. Digestion was performed overnight at 37°C. Peptides were eluted, pooled, and dried using a SpeedVac concentrator. Dried peptides were reconstituted in 60 μL Pierce™ Peptide Retention Time Calibration Mixture (PRTC) buffer (2% acetonitrile, 0.1% formic acid, 25 nM PRTC, pH ≤ 2) prior to LC–MS analysis.

### LC–MS/MS-based proteomics analysis

Peptides were separated on a homemade reversed-phase C18 analytical column coupled to a Thermo Scientific Vanquish™ Neo UHPLC system. Mobile phase A consisted of water containing 0.1% formic acid, and mobile phase B consisted of 80% acetonitrile containing 0.1% formic acid. The gradient was comprised of 4% B for 0.5 min at a constant flow rate of 0.9 μL/min, followed by 4–8% B for 0.1 min at a decreased flow rate of 0.4 μL/min, 8–25% B over 12.3 min, 25–45% B over 6.8 min, 45–55% B over 1.4 min at an increased flow rate of 0.8 μL/min, and 55–99% B over 0.5 min at an increased flow rate of 0.9 μL/min, followed by a 1.4 min hold at 99% B. Peptides were analyzed using an Orbitrap Astral mass spectrometer (Thermo Fisher Scientific) equipped with a nano-electrospray ionization (NSI) source operated at 2.0 kV. Full MS scans were acquired over an *m/z* range of 380–980 in the Orbitrap analyzer at a resolution of 240,000 with an automatic gain control (AGC) target of 500% and a maximum injection time of 3 ms. Data-independent acquisition (DIA) MS/MS spectra were acquired over an *m/z* range of 150–2000 using 2 *m/z* isolation windows, a collision energy (CE) of 25%, an AGC target of 500%, and a maximum injection time of 3 ms in the Astral analyzer. All proteomics data have been deposited to the ProteomeXchange Consortium via the PRIDE partner repository with the dataset identifier PXD080132.

### Data analysis of proteomics

Raw DIA-MS data were processed by PTM Bio (Hangzhou Jingjie Biotechnology Co., Ltd.) using DIA-NN (v1.9) for protein identification and quantification. Protein abundance matrices generated from the DIA-MS analysis were used for downstream analyses. To identify proteins whose abundance changes in *Tmem135^FUN025/FUN025^* mice were restored toward WT levels by fish oil treatment, Pearson correlation analysis was performed using a predefined pattern vector (0, 0, 0, 1, 1, 1, 0, 0, 0), corresponding to WT, WT, WT, Mut, Mut, Mut, FO, FO, and FO samples, respectively. Prior to analysis, missing values were imputed using k-nearest neighbors imputation. For each protein, Pearson correlation coefficients and associated P values were calculated between the imputed abundance profile and the pattern vector. Multiple testing correction was performed using the Benjamini–Hochberg procedure, and proteins with significant correlations were defined as pattern-associated proteins. Proteins showing significant correlation with this pattern were classified as fish oil-responsive proteins. Proteins altered in *Tmem135^FUN025/FUN025^* mice but not restored by fish oil treatment were classified as fish oil-nonresponsive proteins. Selected protein groups, including peroxisomal and mitochondrial proteins, were visualized as heatmaps in R using normalized protein abundance values. Functional enrichment analyses were performed using Kyoto Encyclopedia of Genes and Genomes (KEGG) pathway over-representation analysis.

### LC–MS/MS-based malonyl-proteomic profiling

Dried S-Trap™ eluates were reconstituted in 1 mL of desalting buffer (1% acetonitrile, 0.1% trifluoroacetic acid [TFA], pH < 2). When necessary, the pH was further adjusted to below 2 using TFA. Samples were subsequently loaded onto a HyperSep™ C18 cartridge (50 mg resin bed), desalted, and eluted according to the manufacturer’s instructions. Eluted peptides were frozen and dried using a SpeedVac concentrator. Dried peptides were resuspended in immunoprecipitation (IP) buffer (100 mM NaCl, 1 mM EDTA, 50 mM Tris-HCl, and 0.5% NP-40, pH 8.0). The peptide solution was incubated overnight at 4°C with pre-washed anti-malonyllysine antibody-conjugated agarose beads (PTM904; PTM Bio, Hangzhou Jingjie Biotechnology Co., Ltd.) on a rotary shaker with gentle agitation. Following incubation, the beads were washed four times with IP buffer and twice with deionized water. Bound malonylated peptides were eluted three times using 0.1% TFA, and the eluates were pooled and vacuum-dried. The dried peptides were subsequently desalted using C18 ZipTips according to the manufacturer’s instructions and subjected to LC–MS/MS analysis. Peptides were loaded onto a homemade reversed-phase C18 analytical column connected to a Thermo Scientific Vanquish™ Neo UHPLC system. Separation was performed using a gradient consisting of 4% mobile phase B (80% acetonitrile, 0.1% formic acid) for 0.5 min at a flow rate of 0.9 μL/min, followed by 4–8% B for 0.1 min at a reduced flow rate of 0.4 μL/min, 8–25% B over 12.3 min, 25–45% B over 6.8 min, 45–55% B over 1.4 min at an increased flow rate of 0.8 μL/min, and 55–99% B over 0.5 min at a flow rate of 0.9 μL/min. The gradient was then maintained at 99% B for an additional 1.4 min. The separated peptides were introduced into an Orbitrap Astral mass spectrometer (Thermo Fisher Scientific) via a NSI source operated at 2.0 kV. Full MS spectra were acquired in the Orbitrap analyzer over an *m/z* range of 380–980 at a resolution of 240,000, with an AGC target of 500% and a maximum injection time of 3 ms. DIA MS/MS spectra were collected over an *m/z* range of 150–2000 using 2 *m/z* isolation windows, a CE of 25%, an AGC target of 500%, and a maximum injection time of 3 ms in the Astral analyzer. The mass spectrometry proteomics data for lysine malonylation have been deposited to the ProteomeXchange Consortium via the PRIDE partner repository with the dataset identifier PXD080132.

### Data analysis of malonylated proteomics

To identify malonylation sites that were increased in *Tmem135^FUN025/FUN025^* mice and restored toward WT levels by fish oil treatment, Pearson correlation analysis was performed using a predefined pattern vector (0, 0, 0, 1, 1, 1, 0, 0, 0), corresponding to WT, WT, WT, Mut, Mut, Mut, FO, FO, and FO samples, respectively. Correlation analyses were performed for both malonylation site abundance and total protein abundance. Sites or proteins lacking P values from the initial differential analysis were excluded prior to correlation analysis. Remaining missing values were imputed using the k-nearest neighbor imputation method. Malonylation sites were classified as protein abundance-independent when the correlation between malonylation abundance and the predefined pattern was significant (FDR < 0.2), whereas the corresponding protein abundance was not significantly correlated with the pattern (FDR > 0.2). Differentially malonylated proteins were subjected to Kyoto Encyclopedia of Genes and Genomes (KEGG) pathway over-representation analysis (ORA) using the NetworkAnalyst platform. Enriched pathways were ranked based on false discovery rate (FDR), and the top enriched pathways were visualized as a dot plot in R (v4.4.2) using ggplot2, where dot size represents the number of proteins associated with each pathway and color indicates pathway significance (−log10 FDR).

### L-lactate measurement

L-lactate levels were quantified using a commercially available colorimetric assay kit (#ab65331, Abcam), which is based on an enzymatic reaction in which lactate is oxidized by lactate dehydrogenase, leading to the formation of a chromogenic product through interaction with the probe. Samples were incubated for 30 min at room temperature with the reaction mixture containing lactate assay buffer, lactate enzyme mix, and lactate substrate mix. Absorbance was then measured at 450 nm using a SpectraMax® Plus 384 Absorbance Plate Reader.

### Quantification and statistical analysis

All quantitative data are presented as mean ± SD or mean ± SEM as indicated in the figure legends. Statistical analyses were performed using GraphPad Prism software (GraphPad Software, San Diego, CA, USA). For experiments involving two genetic factors, statistical significance was assessed using two-way analysis of variance (ANOVA) followed by Šídák’s multiple-comparisons test. For comparisons among WT, *Tmem135^FUN025/FUN025^*, and fish oil-fed *Tmem135^FUN025/FUN025^* mice, statistical significance was determined using one-way ANOVA followed by Dunnett’s multiple-comparisons test, with WT serving as the control group. The statistical test used for each dataset is specified in the corresponding figure legends. Statistical significance was defined as *p < 0.05, **p < 0.01, ***p < 0.001, and ****p < 0.0001.

## Notes

### Competing Interest Statement

The authors have declared no competing interest.

