## Supplementary material for "TMEM135 deficiency remodels hepatic lipid homeostasis and protein malonylation through a DHA-sensitive lipogenic program buffered by peroxisomal metabolism": Supplemenary figures

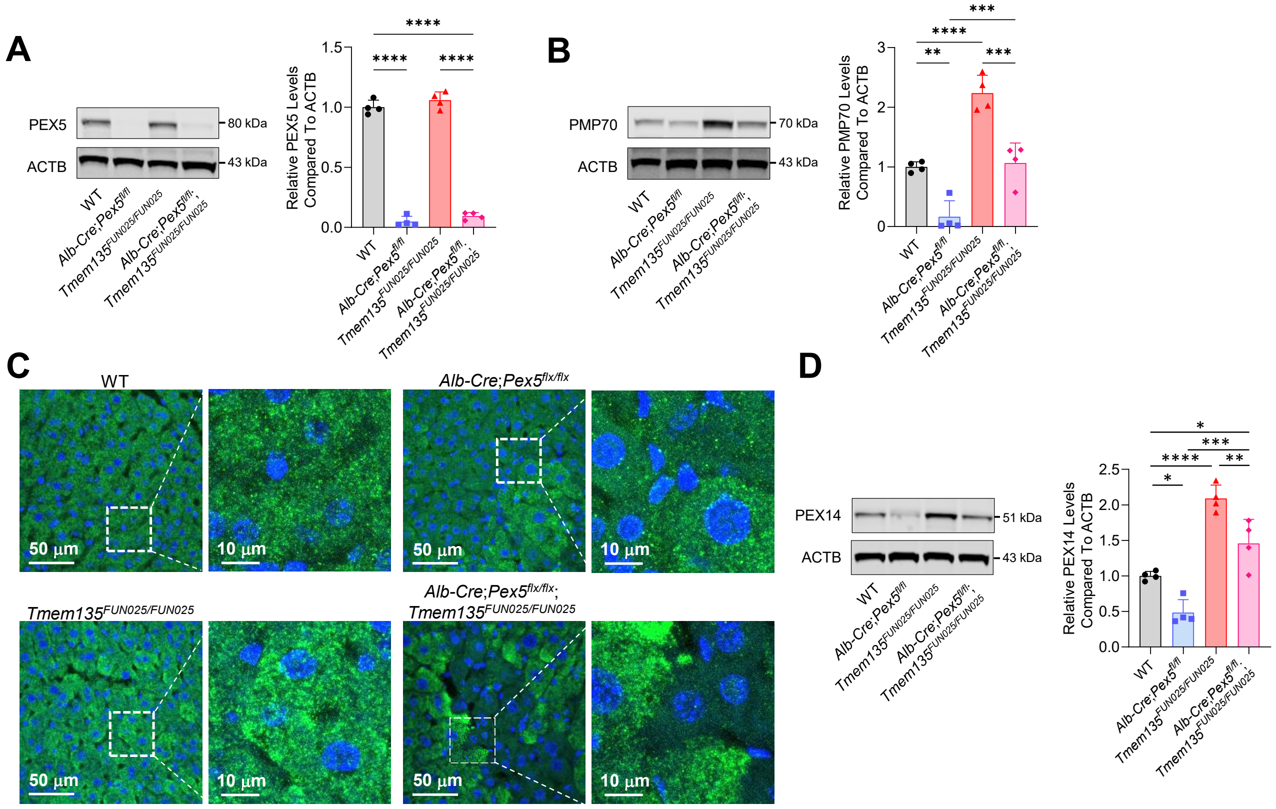


**Figure S1. Validation of liver-specific Pex5 deletion and peroxisome depletion, related to Figure 1.**

(A) Representative immunoblots and quantification of peroxisomal biogenesis factor 5 (PEX5) in the liver lysates from WT, *Alb-Cre*;*Pex5*$^{fl/fl}$, *Tmem135*$^{FUN025/FUN025}$, and *Alb-Cre*;*Pex5*$^{fl/fl}$;*Tmem135*$^{FUN025/FUN025}$ mice.

(B) Representative immunoblots and quantification of 70-kDa peroxisomal membrane protein (PMP70) in the liver lysates from the indicated genotypes.

(C) Representative confocal micrographs of peroxisomal biogenesis factor 14 (PEX14) immunostaining (green) and DAPI (blue) in liver sections from the indicated genotypes. Scale bars, 50 µm (main images) and 10 µm (enlarged images).

(D) Representative immunoblots and quantification of PEX14 in the liver lysates from the indicated genotypes.

All immunoblot quantification data are presented as mean ± SD (n = 4 mice per group). β-actin (ACTB) served as the loading control for these Western blot experiments. Statistical significance was determined by two-way ANOVA followed by Šídák's multiple-comparisons test. *$p < 0.05$, **$p < 0.01$, ***$p < 0.001$, ****$p < 0.0001$

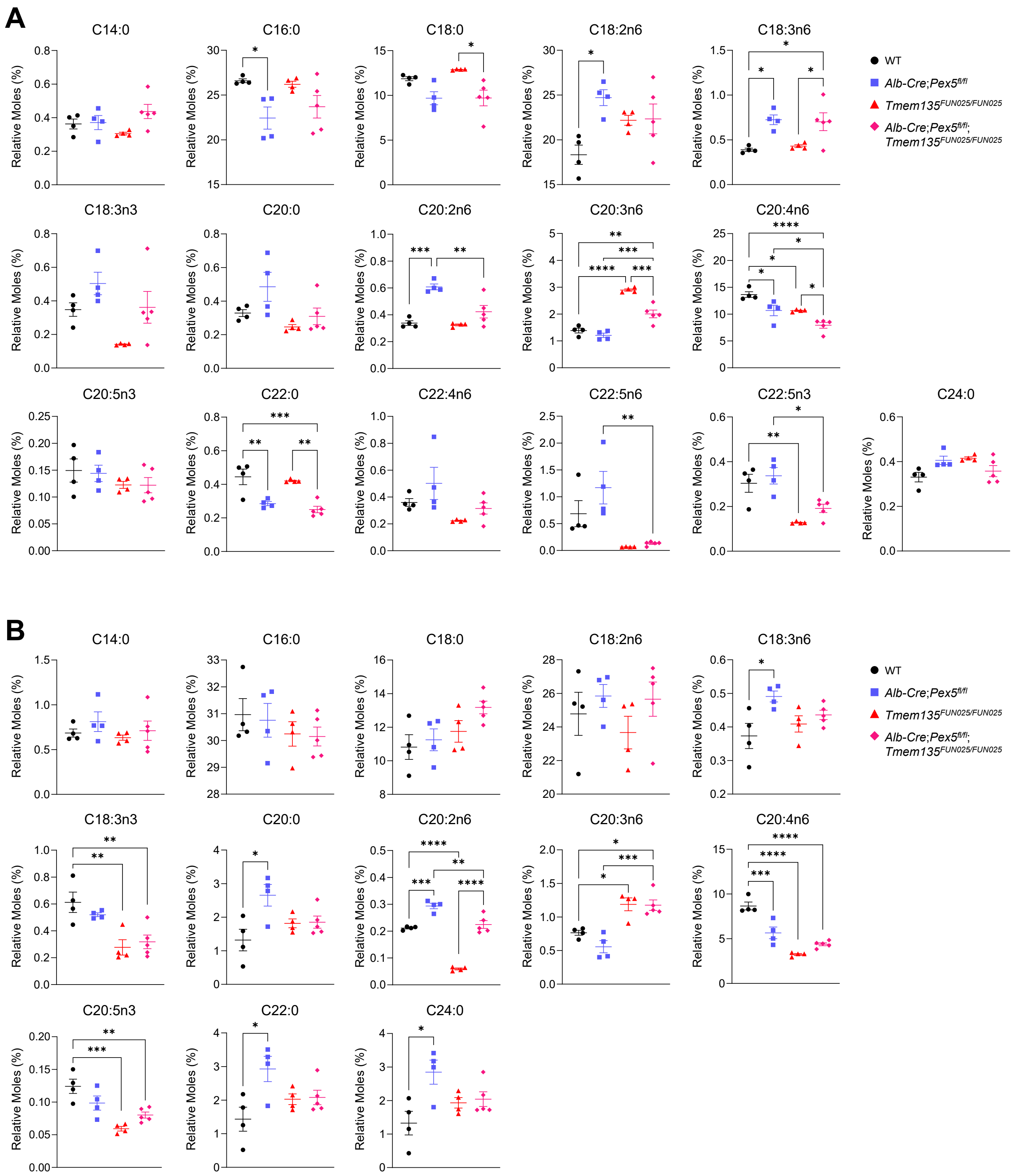


**Figure S2. Detailed fatty acid profiling in liver and plasma across genotypes, related to Figure 2.**

(A) Quantification of hepatic fatty acid levels in WT, *Alb-Cre*;*Pex5*$^{fl/fl}$, *Tmem135*$^{FUN025/FUN025}$, and *Alb-Cre*;*Pex5*$^{fl/fl}$;*Tmem135*$^{FUN025/FUN025}$ mice.

(B) Quantification of plasma fatty acid levels in WT, *Alb-Cre*;*Pex5*$^{fl/fl}$, *Tmem135*$^{FUN025/FUN025}$, and *Alb-Cre*;*Pex5*$^{fl/fl}$;*Tmem135*$^{FUN025/FUN025}$ mice.

Fatty acid quantification data are presented as mean ± SEM (n = 4–5 mice per group). Statistical significance was determined by two-way ANOVA followed by Šídák's multiple-comparisons test. *$p < 0.05$, **$p < 0.01$, ***$p < 0.001$, ****$p < 0.0001$

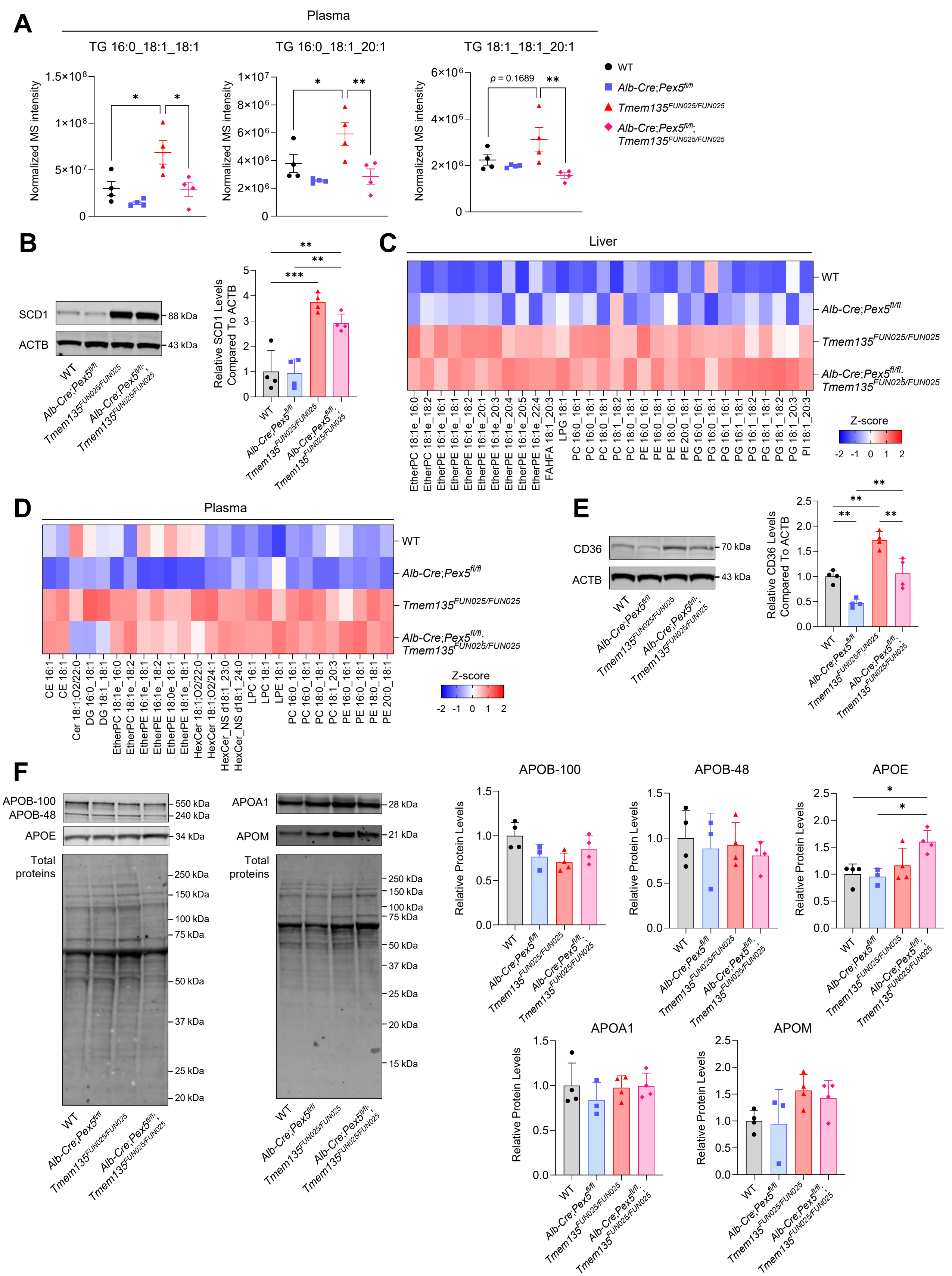

A
Plasma
TG 16:0_18:1_18:1
TG 16:0_18:1_20:1
TG 18:1_18:1_20:1
Normalized MS intensity
p = 0.1689
WT
Alb-Cre;Pex5fl/fl
Tmem135FUN025/FUN025
Alb-Cre;Pex5fl/fl; Tmem135FUN025/FUN025
B
SCD1
ACTB
88 kDa
43 kDa
Relative SCD1 Levels Compared To ACTB
C
Liver
Z-score
D
Plasma
E
CD36
70 kDa
Relative CD36 Levels Compared To ACTB
F
APOB-100
APOB-48
APOE
Total proteins
APOA1
APOM
Relative Protein Levels

**Figure S3. Additional characterization of MUFA-containing lipid species and lipid metabolic regulators, related to Figure 2.**

(A) Quantification of plasma triglyceride (TG 16:0_18:1_18:1, TG 16:0_18:1_20:1, TG 18:1_18:1_20:1) levels in WT, *Alb-Cre*;*Pex5$^{fl/fl}$*, *Tmem135$^{FUN025/FUN025}$*, and *Alb-Cre*;*Pex5$^{fl/fl}$*;*Tmem135$^{FUN025/FUN025}$* mice.

(B) Representative immunoblots and quantification of stearoyl-CoA desaturase 1 (SCD1) in the liver lysates from the indicated genotypes.

(C) Heatmap showing Z-score-normalized abundance of MUFA-containing lipid species in livers from the indicated genotypes.

(D) Heatmap showing Z-score-normalized abundance of MUFA-containing lipid species in plasma from the indicated genotypes.

(E) Representative immunoblots and quantification of CD36 in the liver lysates from the indicated genotypes.

(F) Representative immunoblots and quantification of apolipoprotein B-100 (APOB-100), apolipoprotein B-48 (APOB-48), apolipoprotein E (APOE), apolipoprotein A1 (APOA1), and apolipoprotein M (APOM) in the plasma lysates isolated from 3-month-old mice of the indicated genotypes.

Immunoblot quantification data are presented as mean ± SD (n = 4 mice per group). β-actin (ACTB) served as a loading control for liver lysates, whereas total protein staining was used as the loading control for plasma samples. Triglyceride quantification data are presented as mean ± SEM (n = 4 mice per group). Statistical significance was determined by two-way ANOVA followed by Šídák's multiple-comparisons test. *$p < 0.05$, **$p < 0.01$, ***$p < 0.001$

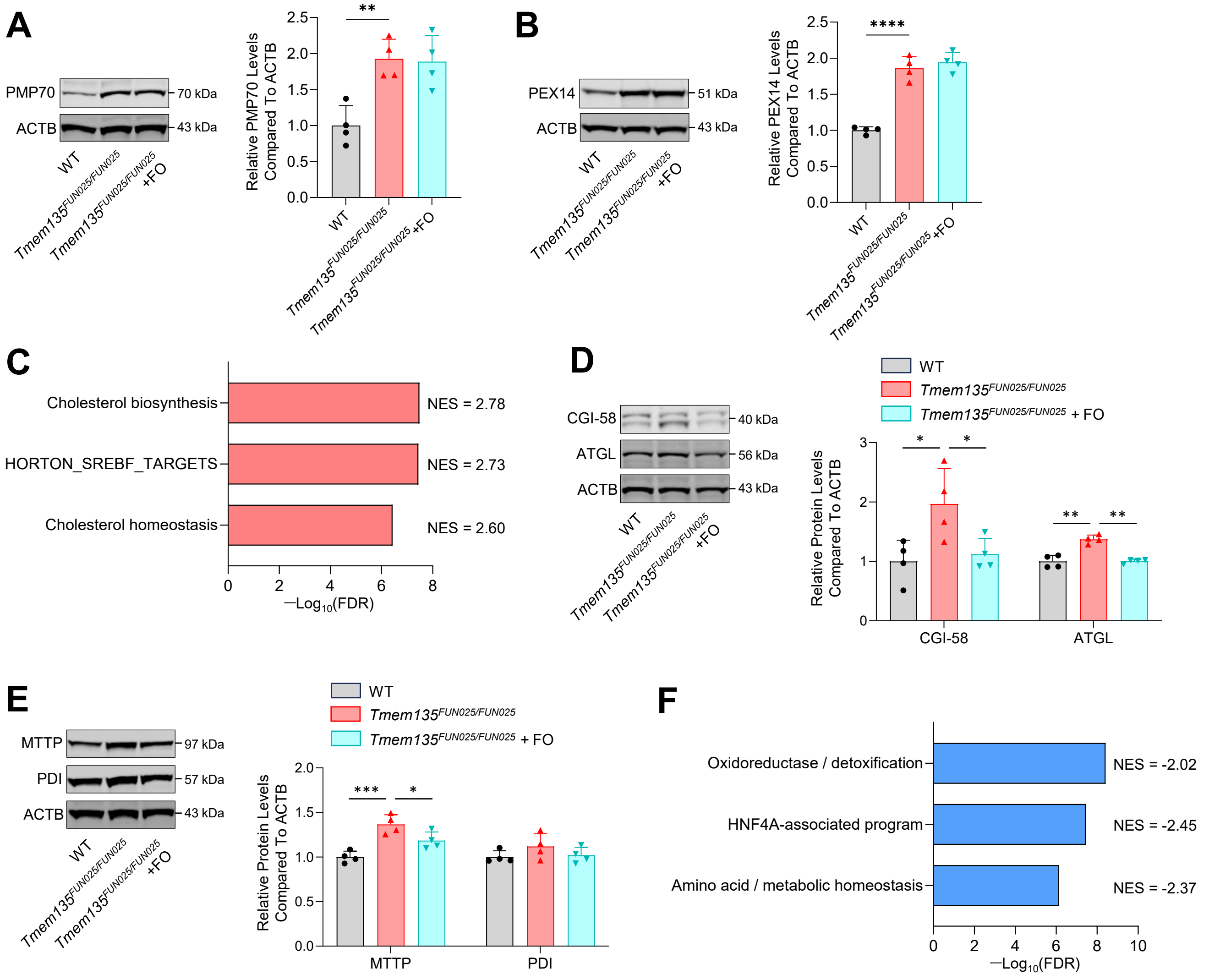


**Figure S4. Validation of metabolic pathways regulated by DHA supplementation in Tmem135 mutant liver, related to Figure 4.**

(A) Representative immunoblots and quantification of 70-kDa peroxisomal membrane protein (PMP70) in the liver lysates from WT, *Tmem135*$^{FUN025/FUN025}$, fish oil (FO)-fed *Tmem135*$^{FUN025/FUN025}$ mice.

(B) Representative immunoblots and quantification of peroxisomal biogenesis factor 14 (PEX14) in the liver lysates from the indicated genotypes.

(C) Gene Set Enrichment Analysis (GSEA) of proteomic data comparing WT and *Tmem135*$^{FUN025/FUN025}$ livers. Lipid biosynthetic and SREBP-dependent metabolic pathways were positively enriched in *Tmem135*$^{FUN025/FUN025}$ livers. NES, normalized enrichment score; FDR, false discovery rate.

(D) Representative immunoblots and quantification of comparative gene identification-58 (CGI-58), and adipose triglyceride lipase (ATGL) in liver lysates from the indicated genotypes.

(E) Representative immunoblots and quantification of microsomal triglyceride transfer protein (MTTP) and protein disulfide isomerase (PDI) in liver lysates from the indicated genotypes.

(F) GSEA of proteomic data comparing WT and *Tmem135*$^{FUN025/FUN025}$ livers. Pathways related to oxidoreductase/detoxification, HNF4A-associated program, and amino acid/metabolic homeostasis were negatively enriched in *Tmem135*$^{FUN025/FUN025}$ livers.
All immunoblot quantification data are presented as mean ± SD (n = 4 mice per group). β-actin (ACTB) served as the loading control for these Western blot experiments. Statistical significance was determined by one-way ANOVA followed by Dunnett's multiple-comparisons test. $^{*}p < 0.05$, $^{**}p < 0.01$, $^{***}p < 0.001$, $^{****}p < 0.0001$

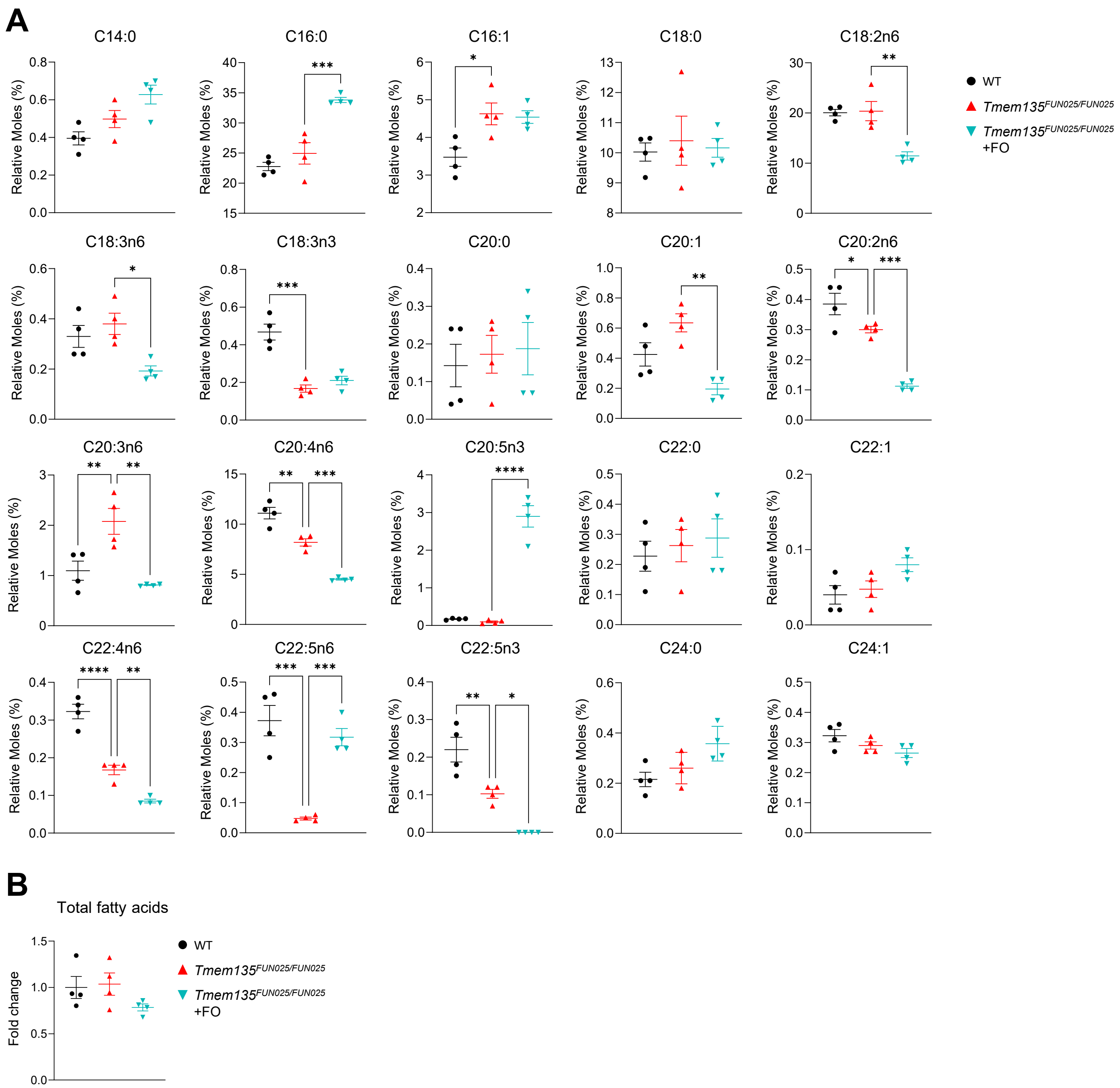


**Figure S5. Hepatic fatty acid profiles following fish oil supplementation, related to Figure 4.**

(A) Quantification of hepatic fatty acid levels in WT, *Tmem135*$^{FUN025/FUN025}$, fish oil (FO)-fed *Tmem135*$^{FUN025/FUN025}$ mice.

(B) Quantification of total hepatic fatty acid content in WT, *Tmem135*$^{FUN025/FUN025}$, FO-fed *Tmem135*$^{FUN025/FUN025}$ mice.

Fatty acid quantification data are presented as mean ± SEM (n = 4 mice per group). Statistical significance was determined by one-way ANOVA followed by Dunnett's multiple-comparisons test. *$p < 0.05$, **$p < 0.01$, ***$p < 0.001$, ****$p < 0.0001$

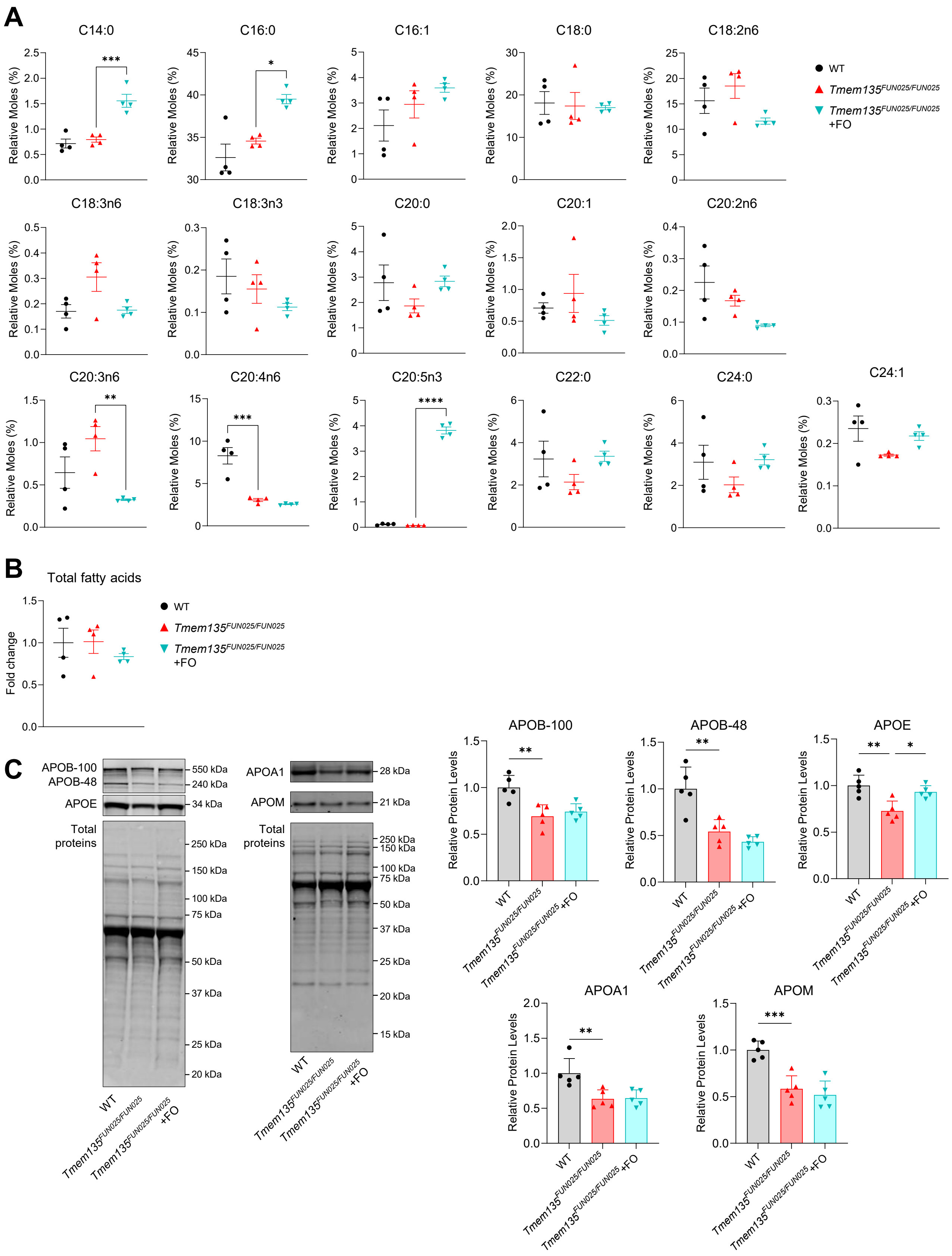
A
C14:0
C16:0
C16:1
C18:0
C18:2n6
C18:3n6
C18:3n3
C20:0
C20:1
C20:2n6
C20:3n6
C20:4n6
C20:5n3
C22:0
C24:0
C24:1
Relative Moles (%)
WT
Tmem135FUN025/FUN025
Tmem135FUN025/FUN025 +FO
B
Total fatty acids
Fold change
C
APOB-100
APOB-48
APOE
Total proteins
APOA1
APOM
550 kDa
240 kDa
34 kDa
28 kDa
21 kDa
250 kDa
150 kDa
100 kDa
75 kDa
50 kDa
37 kDa
25 kDa
20 kDa
15 kDa
Relative Protein Levels

**Figure S6. Effects of fish oil supplementation on plasma fatty acids and apolipoproteins, related to Figure4.**

(A) Quantification of plasma fatty acid levels in WT, *Tmem135*$^{FUN025/FUN025}$, fish oil (FO)-fed *Tmem135*$^{FUN025/FUN025}$ mice.

(B) Quantification of total plasma fatty acid content in the indicated genotypes.

(C) Representative immunoblots and quantification of apolipoprotein B-100 (APOB-100), apolipoprotein B-48 (APOB-48), apolipoprotein E (APOE), apolipoprotein A1 (APOA1), and apolipoprotein M (APOM) in the plasma lysates isolated from 4-month-old mice of the indicated genotypes.

Fatty acid quantification data are presented as mean ± SEM (n = 4 mice per group). Immunoblot quantification data are presented as mean ± SD (n = 4 mice per group). Total protein staining was used as the loading control for plasma samples. Statistical significance was determined by one-way ANOVA followed by Dunnett's multiple-comparisons test. *$p < 0.05$, **$p < 0.01$, ***$p < 0.001$, ****$p < 0.0001$

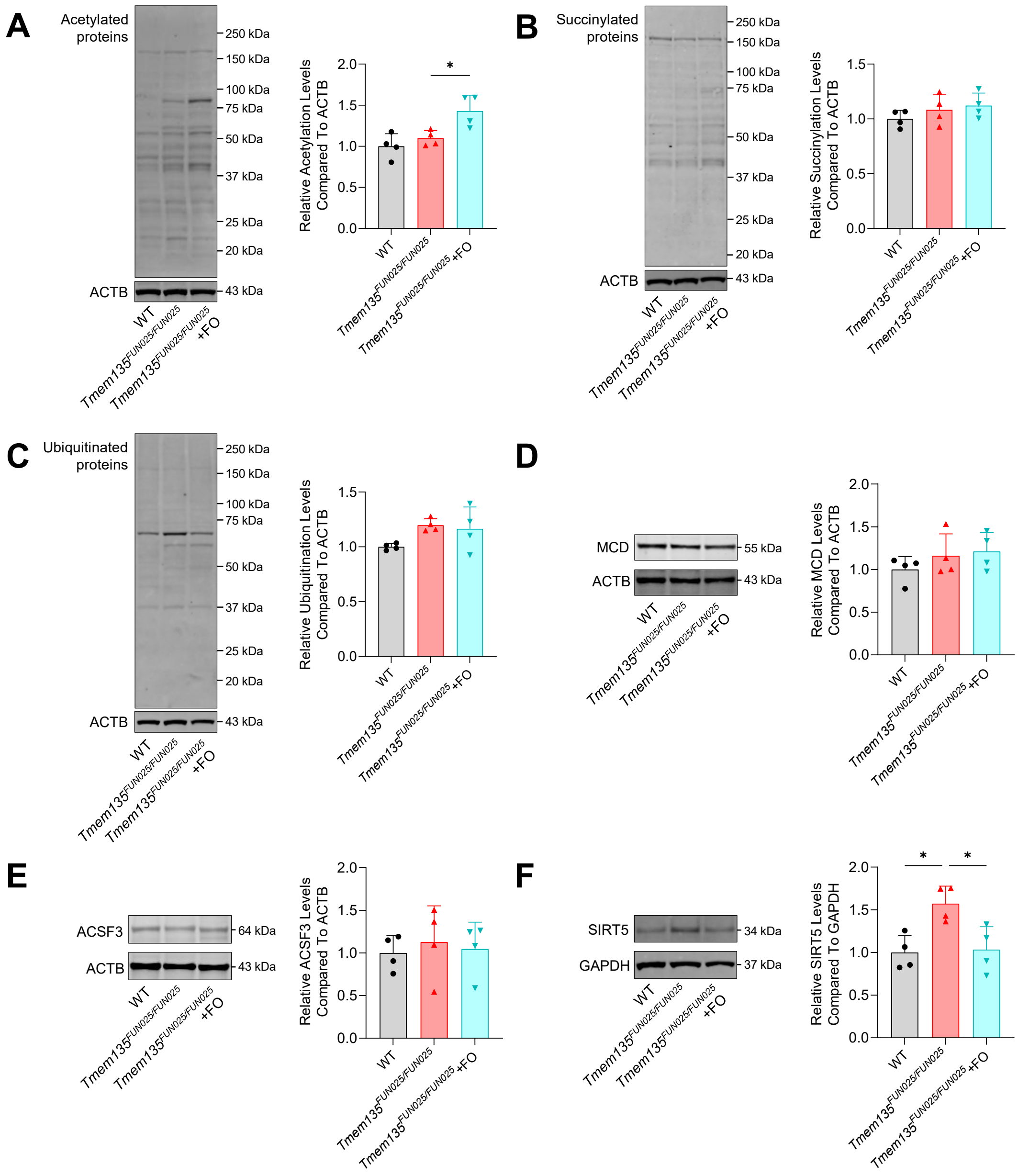

**Figure S7. Selective alteration of protein malonylation in Tmem135 mutant liver, related to Figure5.**

(A) Representative immunoblots and quantification of acetylated proteins in the liver lysates from WT, *Tmem135*$^{FUN025/FUN025}$, fish oil (FO)-fed *Tmem135*$^{FUN025/FUN025}$ mice.

(B) Representative immunoblots and quantification of succinylated proteins in the liver lysates from the indicated genotypes.

(C) Representative immunoblots and quantification of ubiquitinated proteins in the liver lysates from the indicated genotypes.

(D) Representative immunoblots and quantification of malonyl-CoA decarboxylase (MCD) in liver lysates from the indicated genotypes.

(E) Representative immunoblots and quantification of acyl-CoA synthetase family member 3 (ACSF3) in liver lysates from the indicated genotypes.

(F) Representative immunoblots and quantification of sirtuin 5 (SIRT5) in liver lysates from the indicated genotypes

All immunoblot quantification data are presented as mean ± SD (n = 4 mice per group). β-actin (ACTB) served as the loading control for these Western blot experiments. Statistical significance was determined by one-way ANOVA followed by Dunnett’s multiple-comparisons test. *$p < 0.05$

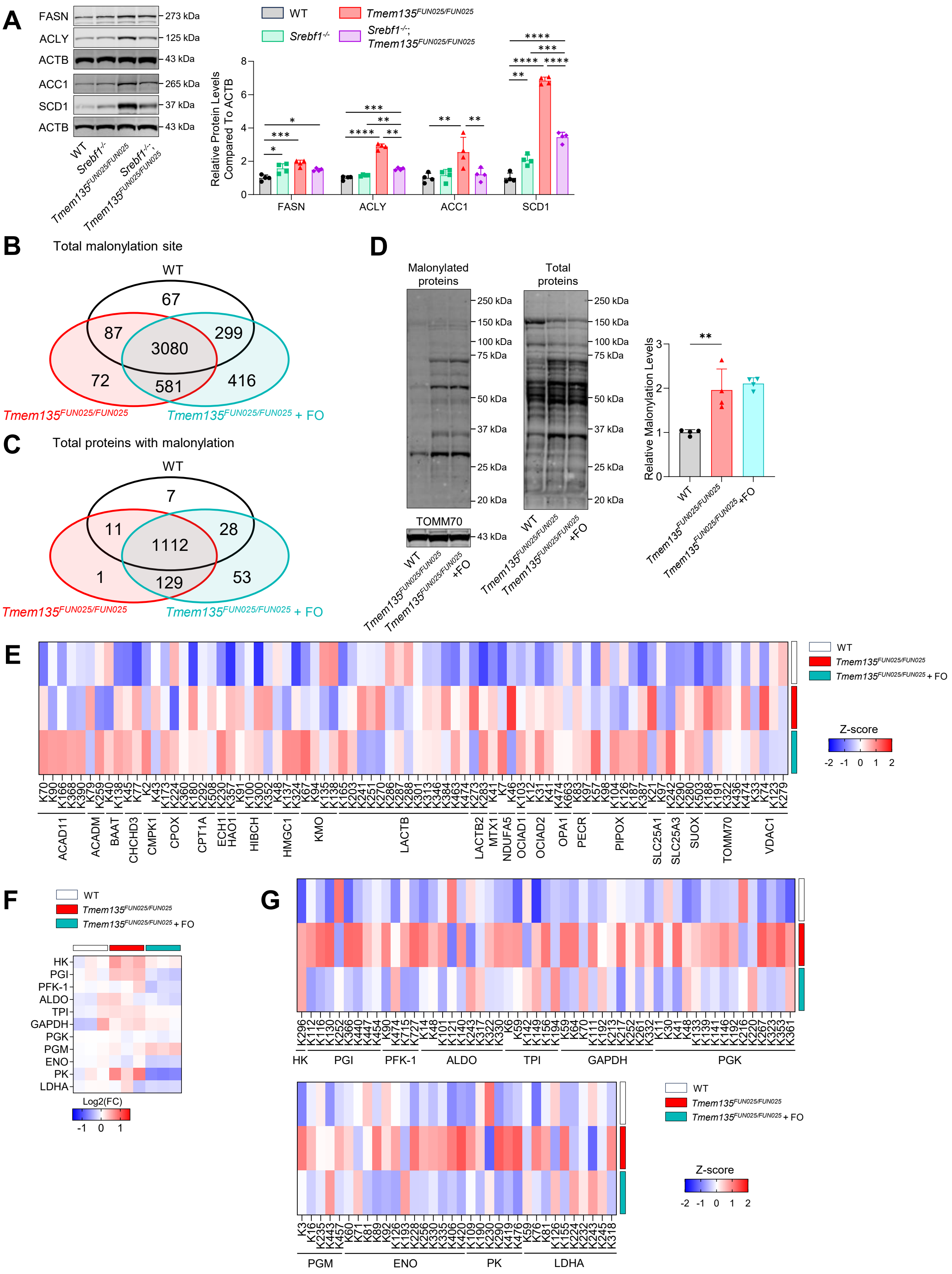
A
FASN
ACLY
ACTB
ACC1
SCD1
ACTB
273 kDa
125 kDa
43 kDa
265 kDa
37 kDa
43 kDa
WT
Srebf1-/-
Tmem135FUN025/FUN025
Srebf1-/-; Tmem135FUN025/FUN025
Tmem135FUN025/FUN025
Srebf1-/-; Tmem135FUN025/FUN025
Relative Protein Levels Compared To ACTB
FASN
ACLY
ACC1
SCD1
B
Total malonylation site
WT
67
87
3080
299
72
581
416
Tmem135FUN025/FUN025
Tmem135FUN025/FUN025 + FO
C
Total proteins with malonylation
WT
7
11
1112
28
1
129
53
D
Malonylated proteins
Total proteins
250 kDa
150 kDa
100 kDa
75 kDa
50 kDa
37 kDa
25 kDa
20 kDa
TOMM70
43 kDa
Relative Malonylation Levels
WT
Tmem135FUN025/FUN025 +FO
E
Z-score
ACAD11
ACADM
BAAT
CHCHD3
CMPK1
CPOX
CPT1A
ECH1
HAO1
HIBCH
HMGC1
KMO
LACTB
LACTB2
MTX1
NDUFA5
OCIAD1
OCIAD2
OPA1
PECR
PIPOX
SLC25A1
SLC25A3
SUOX
TOMM70
VDAC1
F
HK
PGI
PFK-1
ALDO
TPI
GAPDH
PGK
PGM
ENO
PK
LDHA
Log2(FC)
G
HK
PGI
PFK-1
ALDO
TPI
GAPDH
PGK
PGM
ENO
PK
LDHA

**Figure S8. Glycolytic proteins are preferential targets of DHA-sensitive malonylation remodeling, related to Figure 5.**

(A) Representative immunoblots and quantification of the lipogenic enzymes fatty acid synthase (FASN), ATP-citrate lyase (ACYL), acetyl-CoA carboxylase 1 (ACC1), and stearoyl-CoA desaturase 1 (SCD1) in liver lysates from WT, *Srebf1*$^{-/-}$, *Tmem135*$^{FUN025/FUN025}$, *Srebf1*$^{-/-}$;*Tmem135*$^{FUN025/FUN025}$ mice.

(B) Overlap of malonyl-lysine sites identified in the liver malonylomes of WT, *Tmem135*$^{FUN025/FUN025}$, fish oil (FO)-fed *Tmem135*$^{FUN025/FUN025}$ mice.

(C) Overlap of malonylated proteins identified in the liver malonylomes of the indicated genotypes.

(D) Representative immunoblots and quantification of malonylated proteins in the mitochondrial fraction of the indicated genotypes.

(E) Heatmap showing Z score-normalized abundance of malonyl-lysine sites on mitochondrial proteins in livers from the indicated genotypes.

(F) Heatmap showing the relative abundance of glycolysis-related proteins in livers from the indicated genotypes.

(G) Heatmap showing Z-score-normalized abundance of malonyl-lysine sites on glycolysis-related proteins in livers from the indicated genotypes.

Immunoblot quantification data are presented as mean ± SD (n = 4 mice per group). β-actin (ACTB) served as a loading control for liver lysates, whereas total protein staining was used as the loading control for mitochondrial fraction samples. Statistical significance was determined by two-way ANOVA followed by Šídák's multiple-comparisons test for comparisons among WT, *Srebf1*$^{-/-}$, *Tmem135*$^{FUN025/FUN025}$, *Srebf1*$^{-/-}$;*Tmem135*$^{FUN025/FUN025}$ mice, and by one-way ANOVA followed by Dunnett's multiple-comparisons test for comparisons among WT, *Tmem135*$^{FUN025/FUN025}$, and FO-fed *Tmem135*$^{FUN025/FUN025}$ mice. *$p < 0.05$, **$p < 0.01$, ***$p < 0.001$, ****$p < 0.0001$

**Table 1. Antibodies used in western blot analyses**

| Primary antibody | Species | Catalog No. | Manufacturer | RRID |
|---|---|---|---|---|
| ACOX1 | Rabbit | 10957-1-AP | Proteintech | AB_2221670 |
| HSD17B4 (DBP) | Rabbit | 15116-1-AP | Proteintech | AB_2119959 |
| SCP2/SCPx | Rabbit | 23006-1-AP | Proteintech | AB_2879197 |
| PLIN2 | Guinea pig | GP40 | PROGEN | AB_2895086 |
| DGAT2 | Rabbit | NBP1-71701 | Novus Biologicals | AB_11011762 |
| ABHD5 (CGI-58) | Rabbit | 12201-1-AP | Proteintech | AB_2220710 |
| ATGL | Rabbit | 2138 | Cell Signaling Technology | AB_2167955 |
| MTTP | Mouse | BDB612022 | BD Biosciences | AB_399417 |
| PDI | Rabbit | 3501 | Cell Signaling Technology | AB_2156433 |
| SREBP1 | Rabbit | ab28481 | Abcam | AB_778069 |
| ACC1 | Rabbit | 3662 | Cell Signaling Technology | AB_2219400 |
| ACLY | Rabbit | 15421-1-AP | Proteintech | AB_2223741 |
| FASN | Rabbit | ab22759 | Abcam | AB_732316 |
| SCD1 | Rabbit | 2794 | Cell Signaling Technology | AB_2183099 |
| Malonyl-lysine | Rabbit | PTM-901 | PTM BIO | AB_2687947 |
| PEX5 | Rabbit | PA5-58716 | Invitrogen | AB_2645411 |
| PMP70 | Rabbit | ab3421 | Abcam | AB_2219901 |
| PEX14 | Rabbit | 10594-1-AP | Proteintech | AB_2252194 |
| CD36 | Rabbit | 15-674 | ProSci | — |
| APOA1 | Goat | 11A-G2b | Academy Biosciences | — |
| APOB | Goat | AB742 | EMD Millipore | AB_92217 |
| APOE | Rabbit | 68587 | Cell Signaling Technology | AB_3094528 |
| APOM | Mouse | sc-365139 | Santa Cruz Biotechnology | AB_10708274 |
| Acetyl-lysine | Rabbit | 9441 | Cell Signaling Technology | AB_331805 |
| UBC | Rabbit | 10457-1-AP | Proteintech | AB_2241301 |
| Succinyl-lysine | Rabbit | PTM-401 | PTM BIO | AB_2687628 |
| MLYCD/MCD | Rabbit | ab234879 | Abcam | — |
| ACSF3 | Rabbit | PA5-25803 | Invitrogen | AB_2543303 |
| SIRT5 | Rabbit | 8782 | Cell Signaling Technology | AB_2716763 |
| TOMM70 | Rabbit | 14528-1-AP | Proteintech | AB_2303727 |
| ACTB | Mouse | ab8226 | Abcam | AB_306371 |
| ACTB | Rabbit | 4970 | Cell Signaling Technology | AB_2223172 |
| **Secondary antibody** | **Species** | **Catalog No.** | **Manufacturer** | **RRID** |
| IRDye® 680RD Donkey anti-Rabbit IgG | Donkey | 926-68073 | LI-COR Biosciences | AB_10954442 |
| IRDye® 800CW Donkey anti-Rabbit IgG | Donkey | 926-32213 | LI-COR Biosciences | AB_621848 |
| IRDye® 800CW Donkey anti-Guinea Pig IgG | Donkey | 925-32411 | LI-COR Biosciences | AB_2814905 |
| IRDye® 680RD Donkey anti-Goat IgG | Donkey | 926-68074 | LI-COR Biosciences | AB_10956736 |
| IRDye® 800CW Goat anti-Mouse IgG1 | Goat | 926-32350 | LI-COR Biosciences | AB_2782997 |
| IRDye® 800CW Goat anti-Mouse IgG2a | Goat | 926-32351 | LI-COR Biosciences | AB_2782998 |
